# HDAC1 acts as tumor suppressor in ALK-positive anaplastic large-cell lymphoma: Implications for HDAC inhibitor therapy

**DOI:** 10.1101/2024.06.03.597085

**Authors:** Maša Zrimšek, Kristina Draganić, Anna Malzer, Verena Doblmayr, Rafael de Freitas e Silva, Sabrina Wohlhaupter, Carlos Uziel Perez Malla, Katarina Mišura, Heinz Fischer, Helga Schachner, Ana-Iris Schiefer, Raheleh Sheibani-Tezerji, Wilfried Ellmeier, Christian Seiser, Gerda Egger

## Abstract

Histone deacetylases (HDACs) play essential roles in T cell development, and several HDAC inhibitors (HDACi) have gained approval for treating peripheral T cell lymphomas. In this study, we investigated the effects of genetic or pharmacological HDAC inhibition on NPM-ALK positive anaplastic large cell lymphoma (ALCL) development to elucidate potential contraindications or indications for the use of HDACi for the treatment of this rare T-cell lymphoma. Short-term systemic pharmacological inhibition of HDACs using the class I-specific HDACi Entinostat in a premalignant ALCL mouse model postponed or even abolished lymphoma development, despite high expression of the NPM-ALK fusion oncogene. To further disentangle the effects of systemic HDAC inhibition from thymocyte intrinsic effects, conditional genetic deletions of highly homologous class I HDAC1 and HDAC2 enzymes were employed. In sharp contrast to the systemic inhibition, T cell-specific deletion of *Hdac1* or *Hdac2* in the ALCL mouse model significantly accelerated NPM-ALK-driven lymphomagenesis, with *Hdac1* loss having a more pronounced effect. Integration of gene expression and chromatin accessibility data revealed that *Hdac1* deletion selectively perturbed cell type specific transcriptional programs, crucial for T cell differentiation and signaling. Moreover, multiple oncogenic signaling pathways, including PDGFRB signaling, were highly upregulated. The accelerated lymphomagenesis primarily depended on the catalytic activity of HDAC1, as the expression of a catalytically inactive HDAC1 protein showed similar effects to the complete knockout. Our findings underscore the tumor-suppressive function of class I HDAC1 and HDAC2 in T cells during ALCL development, however systemic pharmacological inhibition of HDACs is still a valid treatment strategy, which could potentially improve current therapeutic outcomes.

## INTRODUCTION

Anaplastic large cell lymphoma (ALCL) is a rare, aggressive non-Hodgkin lymphoma of T cell origin, characterized by large pleomorphic and anaplastic lymphoid cells expressing the CD30 antigen. About 60-80% of ALCL cases harbor a characteristic translocation t(2;5)(p23;q35), resulting in a fusion between the anaplastic lymphoma kinase (*ALK*) and nucleophosmin (*NPM1*) genes^1^. The NPM-ALK fusion oncoprotein is a constitutively active kinase that induces a multitude of downstream signaling pathways ultimately driving malignant transformation of T cells and disease progression^2–4^.

Current treatment modalities include poly-chemotherapy (CHOP and CHOP-like regimens) and sometimes radiation as front-line therapy. In refractory or relapsed disease, brentuximab vedotin, a CD30 antibody drug conjugate, may be added, if not previously used in the front-line setting^5^. Moreover, in 2021 the tyrosine kinase inhibitor (TKI) Crizotinib was approved for the treatment of ALK positive (ALK+) ALCL^6^. However, therapy resistance remains a frequent challenge^7–10^, necessitating the exploration of novel treatment strategies. Likewise, non-chemotherapy approaches could mitigate toxicity and reduce the risk of tumor recurrence. HDACi, such as the FDA-approved Belinostat or Romidepsin have been proposed as options for treating relapsed disease^11^.

HDACs are epigenetic enzymes that regulate gene expression by catalyzing the removal of acetyl groups from histones. They are frequently deregulated in hematological malignancies, including leukemias and lymphomas^12^. HDACs can modulate the transcription of oncogenes and tumor suppressor genes, and some HDACs function as the catalytic subunits of multi-protein corepressor complexes, being aberrantly recruited to target genes to drive tumorigenesis^13^. In ALCL, the proapoptotic gene BIM can be epigenetically silenced through the recruitment of the SIN3a corepressor complex, where HDAC1/2 act as catalytic core^14^. In addition to histone proteins, HDACs can deacetylate non-histone proteins and signaling proteins^15^, like STAT3, which is a key signal transmitter in ALCL^4,16^.

Maintaining adequate levels of HDAC1/2 is crucial for normal T cell development, as they are indispensable for preserving the integrity of CD4 lineage T cells by inhibiting RUNX3-CBFβ complexes that can induce CD8 lineage programs in CD4+ T cells^17^. Likewise, dual inactivation of HDAC1/2 in T cells using *Lck-*Cre leads to a developmental blockade, while reduced HDAC activity results in genomic instability and neoplastic transformation^18,19^. Thus, HDAC1/2 exert an essential role in maintaining genome stability and the development of mature T cell populations. Consequently, the use of HDAC inhibitors could potentially accelerate lymphomagenesis, especially under certain (pre-malignant) conditions, as demonstrated in a mouse model of acute promyelocytic leukemia^20^.

To harness the full therapeutic potential of HDAC inhibitors, a comprehensive understanding of HDAC function *in vivo* is essential. Currently, the role of HDACs as transcriptional regulators and the consequences of their perturbations on chromatin and the transcriptome in ALCL are poorly understood. Here we used a murine model of ALCL, driven by the expression of the human fusion oncogene NPM-ALK in T cells, to assess the effects of pharmacological inhibition or genetic deletion of specific class I HDAC isoforms. Systemic administration of HDACi delayed or completely abrogated tumor development, whereas T cell specific depletion of HDAC1 or HDAC2 or inactivation of the catalytic activity of HDAC1 significantly accelerated lymphomagenesis.

## METHODS

### Human samples

All human samples were obtained from the tissue archive of the Department of Pathology with informed written consent following approval by the ethic boards of the Medical University of Vienna (1224/2014). IHC staining was performed and intensity of staining was determined by a trained pathologist. Antibodies are listed in Supplementary Methods.

### Mice

*Cd4*-NPM-ALK transgenic mice^21^, mice carrying loxP-flanked *Hdac1*^22^ or *Hdac2*^22^ and *Cd4*-Cre mice^23^ were crossed to obtain NPM-ALK *Hdac1*KO and NPM-ALK *Hdac2*KO mice. Similarly, NPM-ALK *Hdac1*KO mice were crossed with mice with a Rosa26 knock-in (KI) construct containing the *Hdac1* gene with a His141→Ala point mutation, which results in expression of catalytically inactive HDAC1^24,25^. Genotyping of mice is described in the Supplementary Methods. The genetic background of mice was mixed (C57Bl/6xSV/129). Mice were kept under specific pathogen-free conditions and the experiments were carried out in agreement with the ethical guidelines of the Medical University of Vienna and the Austrian Federal Ministry for Science and Research (BMWF; GZ.: 66.009/0304.WF/V/3b/2014).

### *In vivo* HDACi treatments

Mice were treated with HDACi for two consecutive weeks on five-days-on-two-days-off schedule. 10 µg/g/day of Entinostat (Selleckchem) was administered *via* intraperitoneal (IP) injection. The drug was diluted in 90% sterile filtered corn oil and 10% DMSO.

### Protein isolation, Western blotting

Snap-frozen tissues were lysed in Hunt buffer and processed for SDS-PAGE and Western blot analysis as previously described^15^. Antibodies and buffers are listed in Supplementary Methods.

### FACS immunophenotyping

Cells were stained for viability and cell surface markers (20 min, RT). Cells were fixed with FoxP3 Transcription Factor Fixation/Permeabilization Solution (eBioscience) and stained for ALK (30 min, 4°C), followed by Alexa Fluor 647 goat anti-mouse IgG antibody (30 min, 4°C). Samples were analyzed with a Cytek Aurora cytometer and quantified using FlowJo^TM^ v10.9.0. Software (BD Life Sciences). The gating strategy is displayed in Supplementary Figure 4. Antibodies used are listed in Supplementary Methods.

### ATAC-seq and RNA-seq sample preparation

Snap frozen tumor tissue was lysed and centrifugated. Supernatants were used to isolate RNA (see below). Nuclei were counted and 50,000 nuclei per sample were resuspended in ATAC-seq Reaction Mix. Reactions were incubated at 37°C and subsequently cleaned up using the MiniElute PCR Purification kit. RNA isolation: supernatants were used to isolate RNA following the QIAgen RNeasy protocol with on-column DNase treatment (Turbo^TM^ DNase, 37°C). More detailed info can be found in Supplementary Methods.

### RNA-seq data analysis

FastQC^26^ and MultiQC^27^ were used for quality check. The reads were mapped to GRCm39 mouse reference genome using STAR aligner^28^. Gene counts were obtained with Htseq-count^29^. DESeq2^30^ was used to perform differential gene expression (DE) analysis. Genes with adj p < 0.05 and |LFC| ≥ 1 were considered significantly differentially expressed.

### ATAC-seq data analysis

Initial quality check was done using FastQC^26^. Bowtie2^31^ was used for mapping reads to the reference genome. Peaks were called with MACS2^32^. FRiP scores were calculated and quality check was performed using MultiQC^27^. Counting reads per peak was accomplished with featureCounts^33^. ChIPseeker^34^ was used for peak annotation.

Differential enrichment analysis was done using edgeR^35^. Motif analysis was done using HOMER software^36^.

### Correlation of RNA-seq and ATAC-seq data

Correlation was done using the software developed by Okonechnikov et al.^37^. Publicly available *Mus musculus* HiC data from the NCBI GEO Database (accession no. GSE105918)^38^ were used. Both RNA- and ATAC-seq were normalized using the TPM+log2 method^39^. The median of all transcript lengths per particular gene was used to calculate transcript length per gene. Consensus peak regions were inferred from ATAC-seq data using DiffBind (v.3.0)^40^. Correlations with p < 0.05 were considered significant.

## RESULTS

### HDAC inhibitor treatment before tumor onset significantly restricts NPM-ALK dependent tumor development in ALCL mouse model

To determine expression levels of HDAC1 and HDAC2 in ALCL, tissue microarrays (TMAs) were assessed by immunohistochemistry (IHC). High levels of HDAC1 and HDAC2 were observed in the vast majority of ALCL cases and were comparable to the levels found in angioimmunoblastic T cell lymphoma (AITL) and peripheral T cell lymphoma (PTCL) cases. Importantly, PTCL exhibits aberrant expression of HDACs^41^ and HDACi are already clinically approved for its treatment^42,43^ (**Figure 1A**). Considering this, ALCL patients might benefit from HDACi therapy as well. However, the critical role of HDAC1 and HDAC2 in the maintenance of genome stability and normal T cell development raises potential cautionary implications regarding the use of HDACi. Moreover, high HDAC1 and HDAC2 levels were likewise observed in untransformed T cells in ALCL TMA cases and were not unique to transformed cells. To further study the effects of HDACi on ALCL development, a mouse model expressing the human oncoprotein NPM-ALK in T cells, which mimics human ALK-positive (ALK+) ALCL, was employed^21^. Young pre-tumorigenic NPM-ALK mice were subjected to treatment with the class I-specific HDACi Entinostat that inhibits HDAC1, HDAC2 and HDAC3. Dose escalation of Entinostat in WT mice (2-week treatment, 5-days-on-2-days-off schedule) with 5, 10, 20 and 50 µg Entinostat/g mouse weight/day demonstrated enzyme inhibitory effects as measured by HDAC activity assays in thymocytes after treatment (**Figure 1B**). Entinostat treatment induced a dose-dependent acute thymic involution, with full recovery two weeks post treatment cessation (**Figure 1C, D**). Due to the toxicity of higher doses, 10µg/g/day was selected for treatment of NPM-ALK mice, initiated at 6 weeks of age for two weeks, to investigate whether HDAC inhibition could postpone or accelerate the NPM-ALK lymphomagenesis. Interestingly, HDACi treatment resulted in a notable delay or even prevention of lymphomagenesis, significantly extending median survival of NPM-ALK transgenic mice from 17.9 to 47.4 weeks following the 2-week long treatment (**Figure 1E**). Interestingly, despite the massive decrease in thymus size during the treatment, there was a lack of apoptotic cells and proliferation of thymocytes remained comparable to untreated counterparts (**Figure 1F**). This might indicate that involution could result from a lack of thymic progenitors from the bone marrow to replenish the thymus. This raises the question, whether the effects of HDACi treatment are a result of HDAC inhibition of the (pre-)tumor cells, the thymic microenvironment, progenitor compartments or combinations thereof.

**FIGURE 1:**
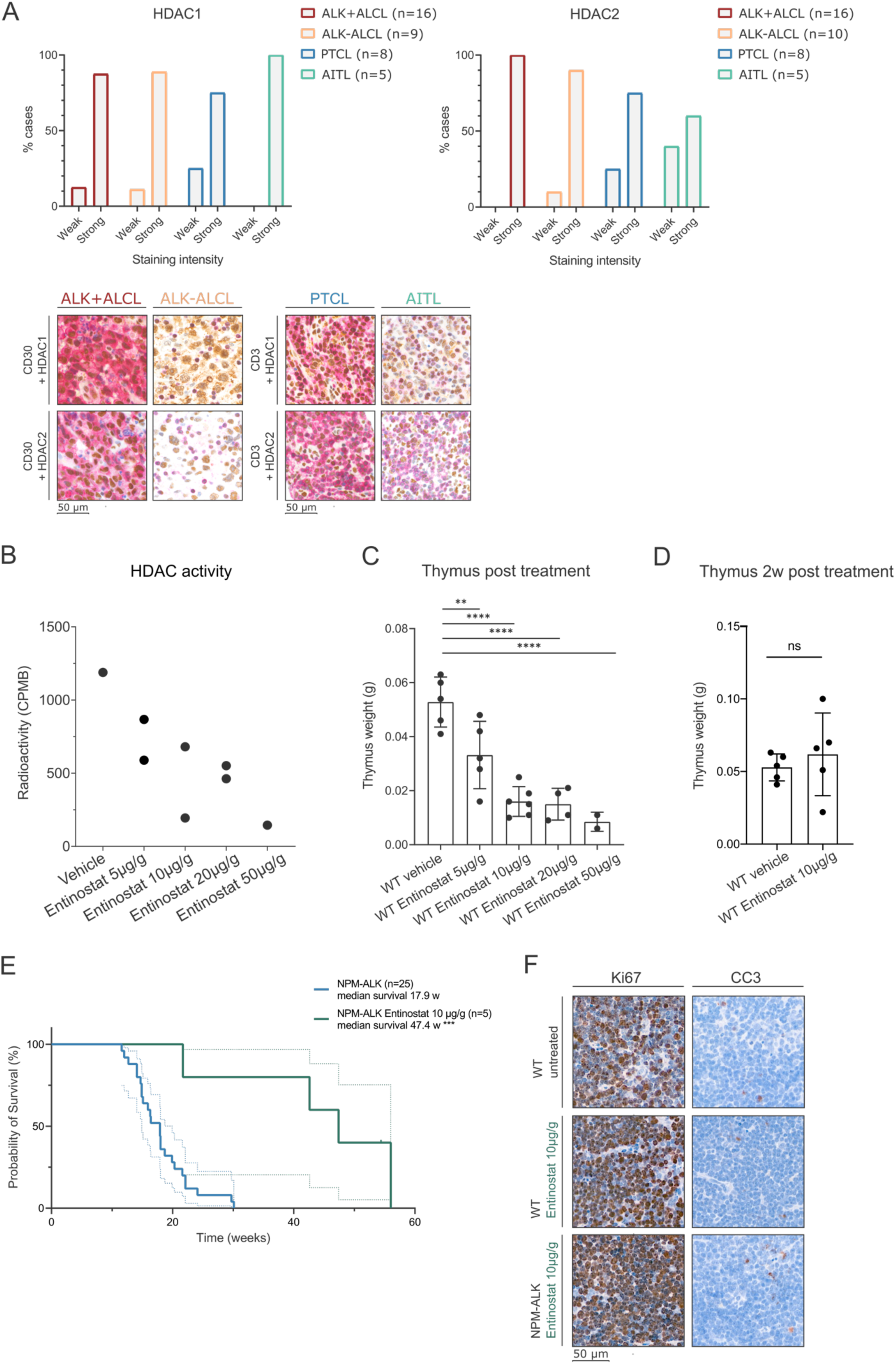
HDAC inhibitor treatment before tumor onset significantly restricts NPM-ALK tumor development in ALCL mouse model. (A) Bar plots depicting the percentages of HDAC1 (left) or HDAC2 (right) staining intensities (weak, strong) on a tissue microarray (TMA) containing specified numbers of ALK+ ALCL, ALK-ALCL, PTCL, and AILT patient samples, evaluated by immunohistochemistry (IHC). The lower panel displays representative microscopic images of IHC stainings from the TMA, as described above (scale bar representing 50 μm). Red cytoplasmic/membrane staining represents CD30/CD3 expression, while brown nuclear staining represents HDAC1/HDAC2 expression. Tissues were stained with the corresponding antibodies and counterstained with hematoxylin (blue). (B) HDAC activity levels in thymi of WT mice after 2-week treatment with Entinostat (n=1 biological replicate for vehicle treatment and 50μg/g/day treatment, n=2 biological replicates for 5μg/g/day, 10μg/g/day and 20μg/g/day treatment, n=2 technical replicates for each biological replicate). Activity levels are measured as counts per minutes beta (CPMB). GraphPad Prism version 8.4.3 was utilized for analysis. (C) Thymic weight of WT mice in biological replicates treated for 2 weeks with vehicle (n=5), 5μg/g/day Entinostat (n=5), 10μg/g/day Entinostat (n=6), 20μg/g/day Entinostat (n=4) and 50μg/g/day Entinostat (n=2). Mean with standard deviation (SD) was plotted using GraphPad Prism version 8.4.3. Statistical significance is indicated by ** for p < 0.01 and **** for p < 0.0001. (D) Thymic weight of WT mice in biological replicates treated for 2 weeks with vehicle (n=5) or 10μg/g/day Entinostat (n=5) and then recovered for an additional 2 weeks. Mean with standard deviation (SD) was plotted using GraphPad Prism version 8.4.3. “ns” indicates not significant. (E) Kaplan Meier survival analysis of NPM-ALK mice (n=25, blue line) and NPM-ALK mice treated with 10μg/g/day Entinostat (n=5, green line) in biological replicates. Median survival of different genotypes was compared using the Log-rank (Mantel Cox) test with GraphPad Prism version 8.4.3. Statistical significance is denoted by *** for p < 0.001. (F) Representative microscopic images of Ki67 and CC3 expression based on IHC staining of thymic sections from untreated WT mice, WT mice treated with 10μg/g/day Entinostat, or NPM-ALK mice treated with 10μg/g/day Entinostat mice. Thymi were excised immediately after the 2-week treatment period. Sections were counterstained with hematoxylin (blue). Scale bar represents 50 μm.

### *Hdac1* loss in T cells accelerates lymphomagenesis

To disentangle the effects of the loss of HDAC activity in pre-tumor cells from the loss in other compartments during ALCL development, *Hdac1* or *Hdac2* were deleted in T cells *via Cd4*Cre recombinase in NPM-ALK mice, resulting in NPM-ALK *Hdac1*KO and NPM-ALK *Hdac2*KO mice (**Figure 2A**). As previously described, NPM-ALK transgenic mice developed thymic tumors with a median survival of 17.9 weeks (**Figure 2B**). Surprisingly, additional deletion of *Hdac1* or *Hdac2* in T cells resulted in strongly accelerated lymphomagenesis with median survivals of 8.1 weeks upon *Hdac1* or 13.75 weeks upon *Hdac2* deletion (**Figure 2B**). Notably, T cell-specific deletion of *Hdac1* induced thymic tumors in approximately a quarter of mice in old age, whereas no signs of malignant transformation were observed in thymi of mice with a deletion of *Hdac2*. (**Supplemental Figure 2A**). This is in line with previous reports demonstrating that the gradual loss of HDAC activity in T cells can result in the induction of thymic lymphomas^18,19^. The size of NPM-ALK and NPM-ALK *Hdac1*KO tumors did not differ, while NPM-ALK *Hdac2*KO mice developed smaller tumors as compared to NPM-ALK mice (**Figure 2C**).

**FIGURE 2:**
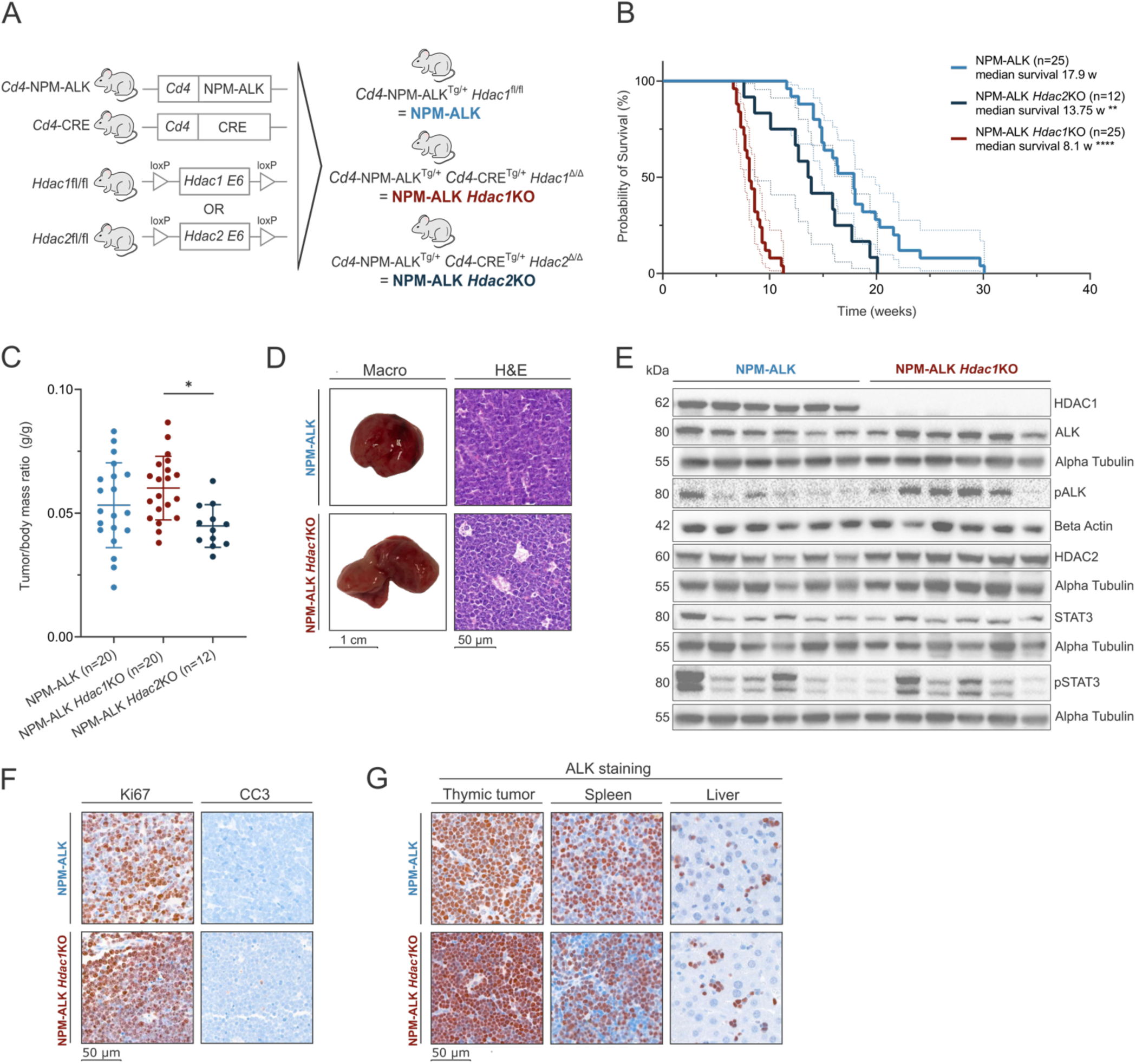
*Hdac1* loss in T cells accelerates lymphomagenesis. (A) Schematic representation of the different mouse models used in the study. The human oncoprotein NPM-ALK was expressed under the T-cell specific *Cd4* promoter. NPM-ALK mice were further crossed with *Cd4*-CRE+ mice with floxed exons 6 of the *Hdac1* or *Hdac2* gene, which produced T cell-specific NPM-ALK transgenic mice with additional *Hdac1* or *Hdac2* knockout (NPM-ALK *Hdac1*KO, NPM-ALK *Hdac2*KO). (B) Kaplan Meier survival analysis of (n=25) NPM-ALK mice (light blue line), (n=12) NPM-ALK *Hdac2*KO mice (dark blue line) and (n=25) NPM-ALK *Hdac1*KO mice (dark red line) in biological replicates. The median survival of different genotypes was compared with Log-rank (Mantel-Cox) test using GraphPad Prism (version 8.4.3). Statistical significance is indicated by ** for p < 0.01 and **** for p < 0.0001. (C) Comparison of thymic tumor(g)/body(g) mass ratios of different genotypes. Mean with standard deviation (SD) is plotted. Groups were compared using one-way Anova corrected for multiple comparison in GraphPad Prism (version 8.4.3). Statistical significance is indicated by * for p < 0.05. (D) Representative macroscopic pictures of end-stage thymic tumors (scale bar representing 1 cm) and hematoxylin-eosin (H&E) stained end-stage thymic tumor sections (scale bar representing 50 μm). (E) Immunoblot of protein expression levels of HDAC1, HDAC2, ALK, pALK, STAT3 and pSTAT3 in end-stage thymic tumors excised from NPM-ALK (n=6) and NPM-ALK *Hdac1*KO mice (n=6). Alpha-Tubulin or Beta-Actin were used as loading controls. The numbers on the left indicate the molecular weight of respective proteins in kiloDalton (kDa). (F) Representative microscopic images of Ki67 and CC3 expression analyzed by IHC of end-stage thymic tumor sections (scale bar representing 50 μm). Ki67 was used as a marker of proliferation and CC3 as a marker of apoptosis. Sections were counterstained with hematoxylin (blue). (G) Representative microscopic images of ALK IHC stainings of end-stage thymic tumor, spleen and liver sections (scale bar representing 50 μm). Sections were counterstained with hematoxylin (blue).

Due to the stronger effects of HDAC1 loss, we further focused on the NPM-ALK *Hdac1KO* model. Differences in macroscopic tissue architecture were observed, with the NPM-ALK tumors mostly being round and encapsulated, while NPM-ALK *Hdac1*KO tumors presented the characteristic two-lobed thymus structure (**Figure 2D**). HDAC1 depletion in NPM-ALK *Hdac1*KO resulted in a compensatory upregulation of HDAC2 protein and induced the activity of the ALK kinase, as indicated by higher levels of phosphorylated ALK (pALK) (**Figure 2E**). Interestingly, pSTAT3, a downstream target of ALK, showed a heterogeneous expression pattern, suggesting that hyperactive ALK signaling upon *Hdac1* deletion might differ from its canonical pathways. Using IHC, no apparent differences in the rate of apoptosis (cleaved caspase 3) or proliferation (Ki67) was observed in end-stage tumors (**Figure 2F**). Nearly 100% of cells in tumors of both genotypes expressed high levels of ALK (**Figure 2G**), and disseminated ALK+ cells were detected in spleen and liver tissues of NPM-ALK and NPM-ALK *Hdac1*KO mice (**Figure 2G**). Especially in the liver, ALK+ cells were prominent around vessels (**Supplemental Figure 2B**), indicating tumor cell dissemination through circulatory and lymphatic systems.

### Accelerated lymphomagenesis depends on HDAC1 enzymatic activity

HDAC1 is part of distinct multi-protein corepressor complexes^44^. Thus, besides its enzymatic function, HDAC1 is also required for complex formation. To disentangle its catalytic and scaffolding functions, we made use of *dHdac1* knock-in (KI) mice that express a catalytically inactive (dead) HDAC1 protein, which can still integrate into corepressor complexes^24,25^. This model reflects the effects of HDACi, which are small molecules binding to the catalytic pocket of HDAC proteins^45^, more closely. The *dHdac1*KI was bred into NPM-ALK *Hdac1*KO mice, generating offspring only expressing catalytically inactive HDAC1 (**Figure 3A**). NPM-ALK *dHdac1*KI mice developed thymic tumors and had a median survival of 9.4 weeks, similar to the complete loss of HDAC1 (8.1 weeks), and again highly accelerated compared to NPM-ALK mice (17.9 weeks) (**Figure 3B, Supplementary Figure 3A**). Interestingly, the *dHdac1*KI was not sufficient to induce thymic lymphomas (**Supplementary Figure 3A**). Tumor sizes were comparable among all genotypes (**Figure 3C**). Same as in NPM-ALK *Hdac1*KO tumors, hyperactivation of the ALK kinase, as indicated by higher levels of phosphorylated ALK (pALK), was observed in NPM-ALK *Hdac1*KI tumors (**Figure D**). The loss of the catalytic HDAC1 activity did not induce a significant upregulation of HDAC2, as previously observed for total HDAC1 depletion, which might explain the significant survival difference of one week between the two genotypes (**Figure 3E, Supplementary Figure 3B**). Further, we performed HDAC activity assays in technical (n=2) and biological (n=5) replicates to measure the overall enzymatic activity of HDACs in tumors of different genotypes. The activity of NPM-ALK *dHdac1*KI samples was significantly lower as compared with NPM-ALK samples (**Figure 3F**). *Hdac1*KO samples showed slightly smaller reduction in overall HDAC activity with borderline significance compared to NPM-ALK tumors (p = 0.059), reflecting the compensatory function of HDAC2. HDAC activity levels of NPM-ALK *Hdac2*KO tumors did not differ from NPM-ALK tumors.

**FIGURE 3:**
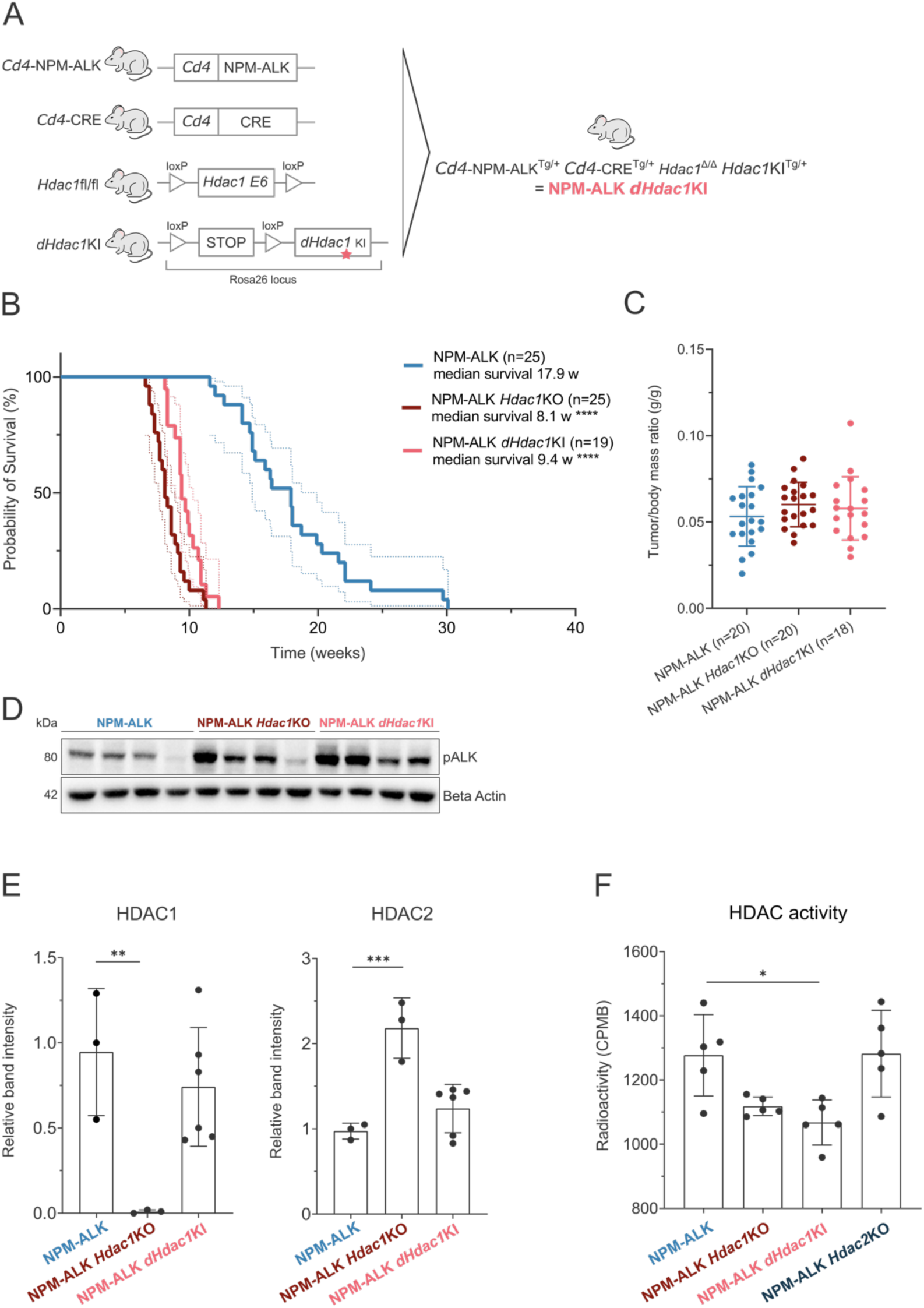
Accelerated lymphomagenesis depends on HDAC1 enzymatic activity. (A) Schematic representation of mouse models used to generate NPM-ALK mice lacking the endogenous *Hdac1* gene but expressing a catalytically dead, mutated HDAC1 protein (dHDAC1) that is unable to deacetylate proteins (NPM-ALK *dHdac1*KI mice). *Cd4* NPM-ALK mice were crossed with CRE+ mice with floxed exons 6 of *Hdac1* to obtain NPM-ALK mice with *Hdac1*KO. These were further crossed with mice containing the *dHdac1* gene inserted into Rosa 26 locus together with a floxed stop cassette. (B) Kaplan Meier survival analysis of NPM-ALK mice (n=25, blue line), NPM-ALK *dHdac1*KI mice (n=19, pink line) and NPM-ALK *Hdac1*KO mice (n=25, red line) in biological replicates. Median survival of different genotypes was compared using the Log-Rank (Mantel-Cox) test with GraphPad Prism (version 8.4.3). Statistical significance is indicated by **** for p < 0.0001. (C) Comparison of thymic tumor(g)/body(g) mass ratios of different genotypes. Mean with standard deviation (SD) is plotted. Groups were compared using one-way Anova corrected for multiple comparison in GraphPad Prism version 8.4.3. (D) Western blot showing protein levels of pALK levels in end-stage thymic tumors excised from NPM-ALK (n=4), NPM-ALK *Hdac1*KO (n=4) and NPM-ALK *Hdac1*KI (n=4) mice. Beta-Actin was used as a loading control. Numbers on the left indicate the molecular weight of analyzed proteins in kiloDalton (kDa). (E) Immunoblot quantification of HDAC1 (left) and HDAC2 (right) protein levels in end-stage thymic tumors of different genotypes. Mean with standard deviation (SD) is plotted. HDAC1 and HDAC2 protein levels were normalized according to Beta-Actin used as a loading control. Groups were compared using one-way Anova corrected for multiple comparison with GraphPad Prism version 8.4.3. Statistical significance is indicated by ** for p < 0.01 and *** for p < 0.001. (F) HDAC activity levels measured in end-stage thymic tumors of different genotypes (n=5 biological replicates, and n=2 technical replicates were used for each genotype). Counts per minutes beta (CPMB) values correspond to HDAC activity levels. Mean with standard deviation (SD) is plotted. Groups were compared using one-way Anova corrected for multiple comparison using GraphPad Prism version 8.4.3. Statistical significance is indicated by * for p < 0.05.

Together, these data suggest that the loss of HDAC1 enzyme activity is the major factor for accelerated lymphomagenesis, and that both HDAC complex formation and HDAC2 activity are less relevant for this process.

### Loss of HDAC1 protein or HDAC1 catalytic activity causes changes in the immunophenotype

To evaluate potential immunophenotypic alterations of HDAC1 depleted tumors, we employed multi-parametric FACS analysis for 19 different immune-cell markers on end-stage tumors, spleens and bone marrow isolated from NPM-ALK, NPM-ALK *Hdac1*KO and NPM-ALK *dHdac1*KI mice. Gating strategies are illustrated in **Supplemental Figure 4A-C**. Unsupervised clustering (tSNE) of CD45^+^ cells showed a clear separation of NPM-ALK tumors from more similar NPM-ALK *Hdac1*KO and NPM-ALK *dHdac1*KI tumors (**Supplemental Figure 4D**). Intracellular staining of tumors for ALK expression revealed that the vast majority of cells was ALK+ for all three genotypes (**Figure 4A, Supplemental Figure 4E**). Interestingly, a higher heterogeneity in the number of ALK+ cells was observed in individual NPM-ALK tumors. The analysis of the thymocyte population based on CD4 and CD8 expression (CD4^−^CD8^−^ double-negative, DN; CD4^+^CD8^+^ double-positive, DP; and CD4^+^CD8^−^ single-positive, CD4SP, CD4^−^CD8^+^ CD8SP) showed that NPM-ALK *Hdac1*KO and NPM-ALK *dHdac1*KI tumors exhibited more cells at the DP stage, while NPM-ALK tumors showed the highest proportion of cells in the DN stage (**Figure 4B**). For the analysis of the CD4^−^CD8^−^ DN population (DN1-4 stages; DN1: CD44^+^CD25^−^; DN2: CD44^+^CD25^+^; DN3: CD44^−^CD25^+^; DN4: CD44^−^CD25^−^), DN cells in all tumors appeared predominantly in the DN4 stage (**Figure 4C**). We further assessed whether ALK+ tumor cells expressing CD4 or CD8 also expressed CD44 or CD62L markers, usually used to distinguish naïve (CD62L^high^ CD44^low^), effector (CD62L^low^ CD44^high^) or central memory T cells (CD62L^high^ CD44^high^). While ALK+ cells from NPM-ALK tumors exhibited considerable heterogeneity, ALK+ tumor cells from NPM-ALK *Hdac1*KO and NPM-ALK *dHdac1*KI tumors mostly resembled a naïve CD4 or CD8 phenotype (CD62L^high^ CD44^low^) (**Figure 4D, 4E**). Consistent with this, the expression of the CD62L, a homing receptor for secondary lymphoid organs, separated NPM-ALK from NPM-ALK *Hdac1*KO and NPM-ALK *dHdac1*KI tumors in the unsupervised clustering analysis (**Supplemental Figure 4F**). NPM-ALK transformed T cells were previously shown to downregulate the TCRb^46^, a finding confirmed in our tumors (**Figure 4F**). Interestingly, ALK+ cells from NPM-ALK *Hdac1*KO and NPM-ALK *dHdac1*KI tumors showed a higher frequency of TCRb+ cells (**Figure 4F**). Since it was hypothesized before that transformed cells require transient TCR expression for thymic egress^47^, we further analyzed ALK+ cells in spleen and bone marrow. We observed a trend of higher ALK+ cell colonization in spleen and bone marrow of NPM-ALK *Hdac1*KO and NPM-ALK *dHdac1*KI compared to NPM-ALK mice (**Figure 4G**). Generally, ALK+ cells found in spleens showed higher frequency of TCRb as compared to their thymic counterparts, which was again more pronounced in tumors lacking HDAC1 protein or enzymatic activity (**Figure 4H**). Together, these data suggest that loss of HDAC1 protein and activity results in a shift of the immunophenotype of ALK+ tumor cells, with higher numbers of cells expressing the TCR, seemingly facilitating increased tumor cell dissemination to distant sites.

**FIGURE 4:**
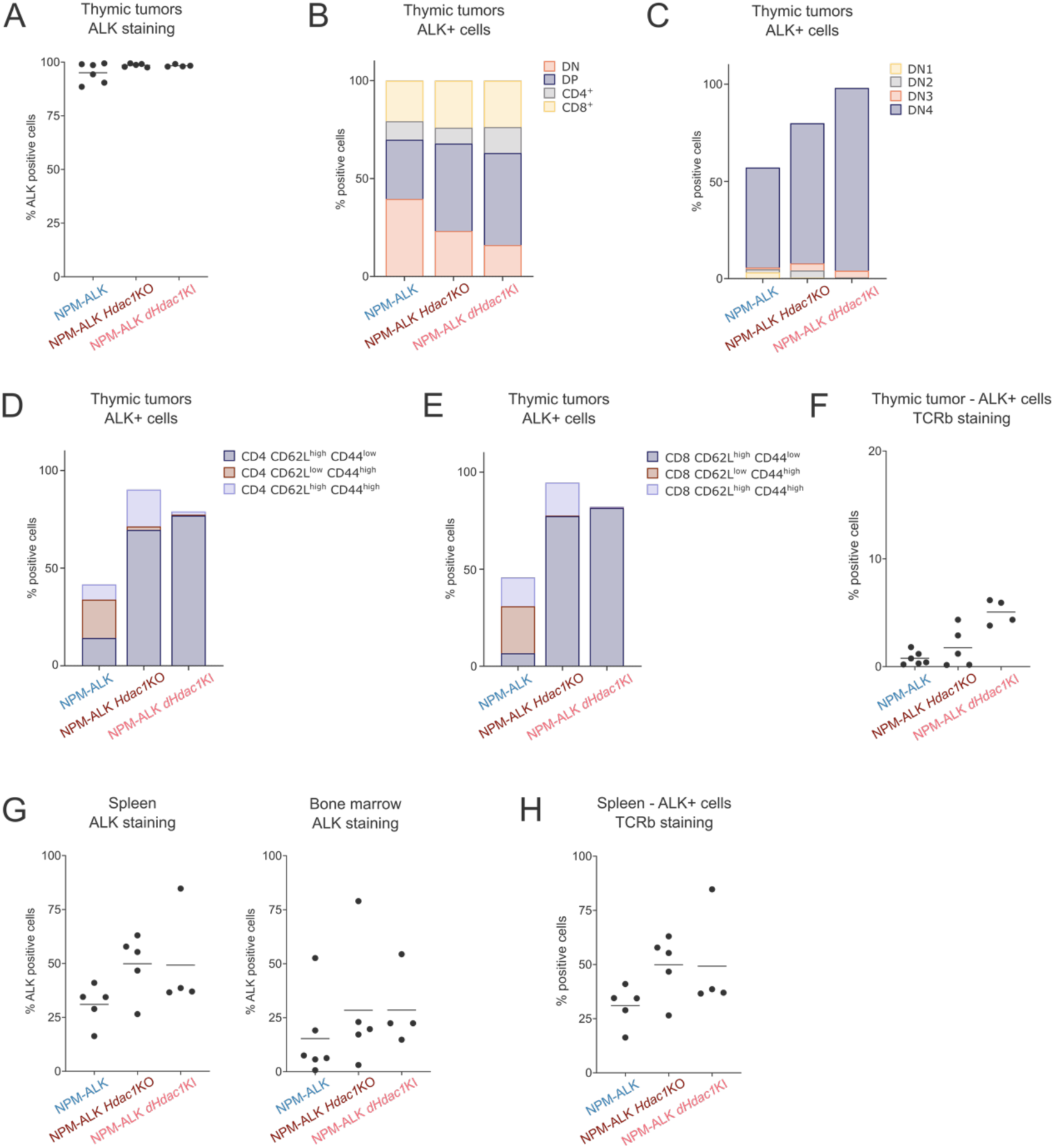
Loss of HDAC1 protein or HDAC1 catalytic activity causes changes in immunophenotype. (A) Percentage of ALK+ cells in end-stage thymic tumors of different genotypes (NPM-ALK, NPM-ALK *Hdac1*KO and NPM-ALK *Hdac1*KI) assessed by FACS. The horizontal line represents the average percentage of positive cells of biological replicates for each genotype. (B) The average distribution of markers for double negative (DN; CD4^−^CD8^−^), double positive (DP; CD4^+^CD8^+^), CD4^+^ and CD8^+^ cells among ALK+ cells in end-stage thymic tumors of indicated genotypes (NPM-ALK, NPM-ALK *Hdac1*KO and NPM-ALK *Hdac1*KI) (C) The average percentage of ALK+ cells exhibiting features of double negative 1 (DN1), double negative 2 (DN2), double negative 3 (DN3), and double negative 4 (DN4) cells in end-stage thymic tumors of different genotypes (NPM-ALK, NPM-ALK *Hdac1*KO and NPM-ALK *Hdac1*KI). (D) The average percentage of ALK+ cells exhibiting features of CD4 naïve (CD4 CD62L^high^ CD44^low^), CD4 effector (CD4 CD62L^low^ CD44^high^) or CD4 central memory T cells (CD4 CD62L^high^ CD44^high^) in end-stage thymic tumors of different genotypes (NPM-ALK, NPM-ALK *Hdac1*KO and NPM-ALK *Hdac1*KI). (E) The average percentage of ALK+ cells exhibiting features of CD8 naïve (CD8 CD62L^high^ CD44^low^), CD8 effector (CD8 CD62L^low^ CD44^high^) or CD8 central memory T cells (CD8 CD62L^high^ CD44^high^) in end-stage thymic tumors of different genotypes (NPM-ALK, NPM-ALK *Hdac1*KO and NPM-ALK *Hdac1*KI). (F) Percentage of ALK+ cells expressing TCRb in end-stage thymic tumors of different genotypes (NPM-ALK, NPM-ALK *Hdac1*KO and NPM-ALK *Hdac1*KI). Average percentage of positive cells of biological replicates of each genotype is represented by the horizontal line. (G) Percentage of ALK+ cells in spleen (left) and bone marrow (right) isolated from mice of different genotypes (NPM-ALK, NPM-ALK *Hdac1*KO and NPM-ALK *Hdac1*KI) presented with end-stage thymic tumors. The horizontal line represents the average percentage of positive cells of biological replicates for each genotype. (H) Percentage of ALK+ cells expressing TCRb in spleen isolated from mice of different genotypes (NPM-ALK, NPM-ALK *Hdac1*KO and NPM-ALK *Hdac1*KI) presented with end-stage thymic tumors. The horizontal line represents the average percentage of positive cells of biological replicates for each genotype.

### Loss of *Hdac1* selectively perturbs cell-type specific transcription

To further evaluate the consequences of *Hdac1* loss on chromatin architecture and gene expression, we focused on end-stage NPM-ALK and NPM-ALK *Hdac1*KO tumors and performed parallel ATAC- and RNA-sequencing on (n=4) biological replicates of both genotypes (**Figure 5A**). Principal component analysis revealed a clear separation of the two groups based on their accessible chromatin regions (**Figure 5B**). Transmission electron microscopy showed similar nuclear structures and chromatin architecture in the different genotypes (**Supplemental Figure 5A**). Notably, *Hdac1* loss did not result in stochastic or global chromatin opening, as the number and size of the peaks representing accessible chromatin were comparable between the two genotypes, with more than 36,000 overlapping peaks (**Figure 5C**). Nevertheless, genotype-specific accessible chromatin regions were identified, encompassing 18,736 in NPM-ALK and 8,613 unique peaks in NPM-ALK *Hdac1*KO tumors (**Figure 5D**). RNA-seq analyses of the same tumors revealed 785 up- and 723 downregulated genes between the two groups (adj p < 0.05 and |LFC| ≥ 1) (**Supplemental Figure 5B**).

**FIGURE 5:**
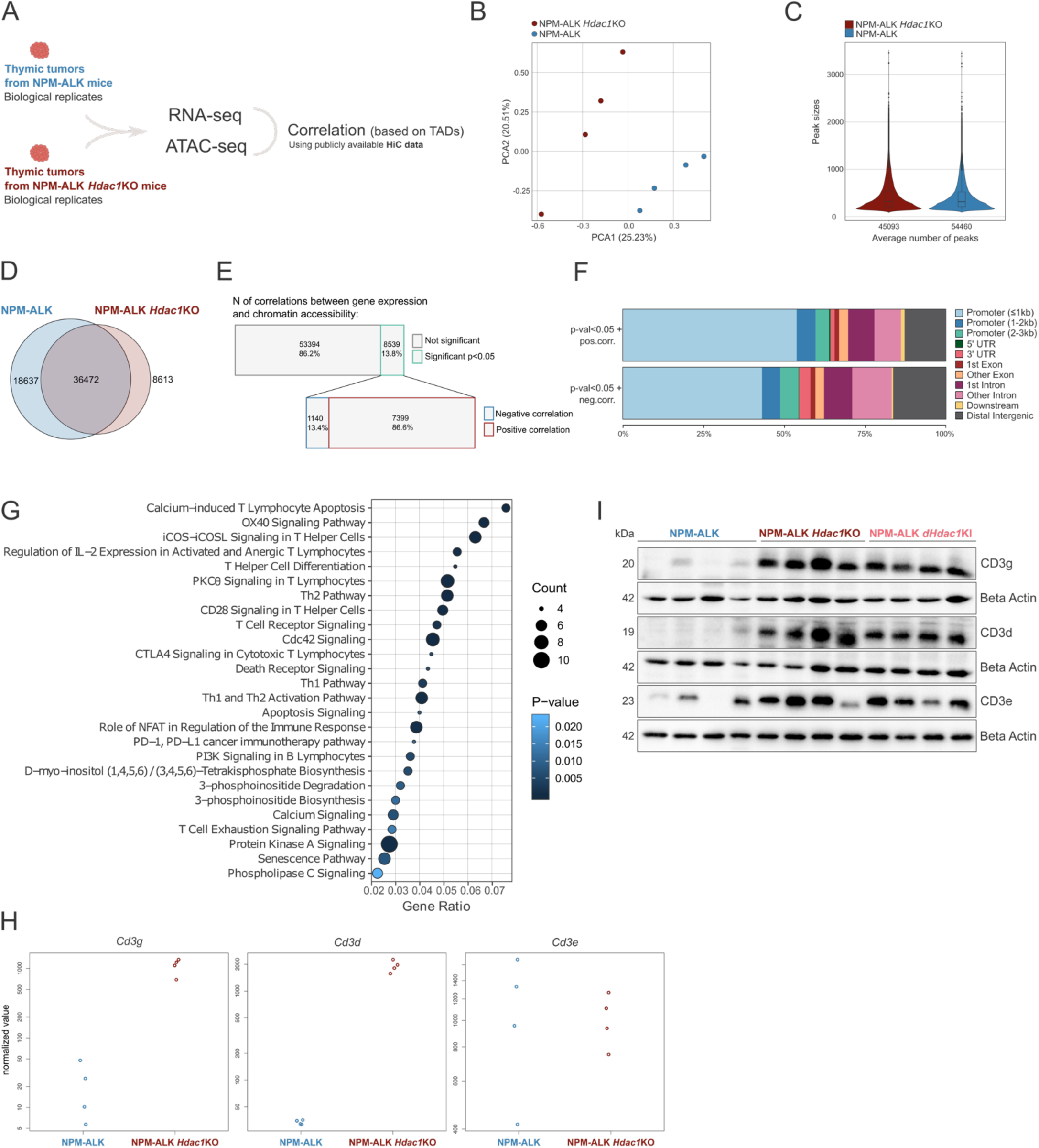
Loss of *Hdac1* selectively perturbs cell-type specific transcription. (A) Schematic representation of ATAC- and RNA-seq experiments. ATAC- and RNA-seq was performed in parallel using end-stage thymic tumors from NPM-ALK mice (biological replicates n=4, blue) and NPM-ALK *Hdac1*KO mice (biological replicates n=4, red). ATAC- and RNA-seq was correlated with topologically associating domains (TADs) inferred from publicly available HiC data^1^. (B) Principal Component Analysis (PCA) of ATAC-seq data illustrating the similarity/variance of NPM-ALK (blue) and NPM-ALK *Hdac1*KO (red) samples. (C) Violin plot showing the distribution of peak sizes and the average number of peaks in NPM-ALK (blue) and NPM-ALK *Hdac1*KO (red) samples based on ATAC-seq analyses. (D) Venn diagram depicting shared and unique open chromatin regions (=peaks) between NPM-ALK (blue) and NPM-ALK *Hdac1*KO (red) samples. (E) Bar charts representing the number and percentages of overall and statistically significant (p < 0.05) correlations between RNA- and ATAC-seq data (upper), as well as the number and percentage of negative and positive correlations among the statistically significant (p < 0.05) correlations (lower). (F) Stacked bar chart representing the percentage of open chromatin regions located in different genomic regions, according to the legend on the right side, comparing the open chromatin regions that are positively correlated (p < 0.05) with changes in gene expression (e.g. open chromatin in promoter region leads to higher gene expression) (upper) and open chromatin regions that are negatively correlated (p < 0.05) with changes in gene expression (e.g. open chromatin in promoter region leads to lower gene expression) (lower). (G) Bubble chart representing significantly enriched pathways in NPM-ALK *Hdac1*KO end-stage thymic tumors as compared to NPM-ALK end-stage thymic tumors based on Ingenuity Pathway Analysis (IPA®) of upregulated genes (RNA-seq: |LFC| ≥ 1, adj p < 0.05) that were correlated with changes in chromatin accessibility (correlation p < 0.5). The size of the circles represents the number of genes affected within a given pathway, the color indicates the significance level based on the gradient scheme on the right. (H) Normalized value for size factor for raw count visualization based on RNA-seq analysis for *Cd3g*, *Cd3d* and *Cd3e* comparing NPM-ALK (blue) and NPM-ALK *Hdac1*KO (red) samples. (I) Immunoblot showing protein levels of CD3d, CD3g and CD3e in end-stage thymic tumors excised from NPM-ALK (n=4), NPM-ALK *Hdac1*KO (n=4) and NPM-ALK *Hdac1*KI mice (n=4). Beta-Actin was used as a loading control. Numbers on the left indicate the molecular weight of analyzed proteins in kiloDalton (kDa).

Next, ATAC-seq data were integrated with RNA-seq data to discern functional promoters/enhancers associated with alterations in chromatin accessibility and dysregulated transcription. It was previously shown that only 47% of distal and proximal elements exhibit interactions with their nearest expressed transcription start site, indicating the necessity to consider long-range promoter/enhancer interactions^48^ and to not use the linear proximity as the only point of reference when correlating the enhancers with their potential target genes. Thus, publicly available chromosome conformation capture (HiC) data of T cells were used to delineate cell type specific topologically associated domains (TADs)^38^. Correlations between open chromatin regions in proximal and distal regulatory elements and transcriptionally active genes were then inferred within the boundaries of every TAD. A total of 8,539 significant correlations were identified (p < 0.05). Among these, 7,399 represented positive correlations, signifying that chromatin opening in promoter/enhancer regions corresponded to higher gene expression, while 1,140 displayed negative correlations, where open chromatin regions corresponded to decreased expression (**Figure 5E**). Peaks associated with up- or downregulated expression did not differ in their size (data not shown), however, positively correlated peaks were more frequently observed in promoter regions as compared to negatively correlated ones, which showed more frequent association with intronic or distal intergenic regions (**Figure 5F**).

Ingenuity Pathway Analysis (IPA®)^49^ of upregulated genes (adj p < 0.05 and |LFC| ≥ 1) with changed chromatin accessibility (correlation p < 0.05) in NPM-ALK *Hdac1*KO tumors revealed that the loss of *Hdac1* selectively perturbed cell type-specific transcription, with top upregulated genes implicated in T cell activation and pathways critical for T cell proliferation and survival (**Figure 5G, Supplemental Table 1**). Of note, increased expression levels of *Cd4* and *Cd8* (Cd4: LFC = 2.27, adj p = 4.90E-04; Cd8: LFC = 3.6, adj p = 2.40E-02) were observed, in line with the increase in DP thymocytes in NPM-ALK *Hdac1*KO tumors seen in immunophenotyping analysis (**Figure 4B**). Furthermore, a marked upregulation of the CD3d/g/e TCR co-receptor was detected in NPM-ALK *Hdac1*KO tumors both on mRNA (**Figure 5H**) and protein levels (**Figure 5I**), consistent with ATAC-seq data showing increased chromatin accessibility in the CD3g/d promoter regions (**Supplemental Figure 5C**). The upregulation of CD3d/g/e was further confirmed in NPM-ALK *Hdac1*KI tumors, mimicking the NPM-ALK *Hdac1*KO tumors (**Figure 5I**).

These findings are in line with previous studies demonstrating the essential function of HATs and HDACs to maintain transcription factor (TF)-dependent lineage-specific gene expression programs^50,51^ and suggest that the T cell receptor (TCR) and its co-receptors, which are usually silenced in ALCL^46,52^, remain active upon HDAC1 depletion.

### Loss of *Hdac1* hyperactivates oncogenic transcription

To delineate the molecular mechanisms driving accelerated lymphomagenesis, we scrutinized chromatin and transcriptional alterations in previously identified HDAC1 and NPM-ALK target genes. The MYC oncogene, a central factor for ALK-driven lymphomagenesis^53,54^, showed comparably high chromatin accessibility in its promoter/enhancer regions as well as mRNA and protein expression in NPM-ALK and NPM-ALK *Hdac1*KO tumors (**Supplemental Figure 6A-C**). NPM-ALK *Hdac1*KO tumors displayed augmented promoter accessibility (LFC = 1.52, p = 1.93E-4) (**Figure 6A**) and gene expression (LFC = 3.76, adj p = 9.5E-10) of the *Jpd2* gene (**Figure 6B**), a MYC-collaborating and p53-suppressing factor previously shown to be upregulated in T cell lymphomas that developed as a consequence of loss of HDAC activity^19^. Furthermore, the oncogenic kinase gene *Tnk2*, implicated in cell survival, proliferation, and migration was upregulated in NPM-ALK *Hdac1*KO tumors (LFC = 2.43, adj p = 2.50E-10) (**Figure 6C**), which potentially enhanced the NPM-ALK oncogenic cascade *via* interaction with NPM-ALK and co-activation of STAT signaling^55^.

**FIGURE 6:**
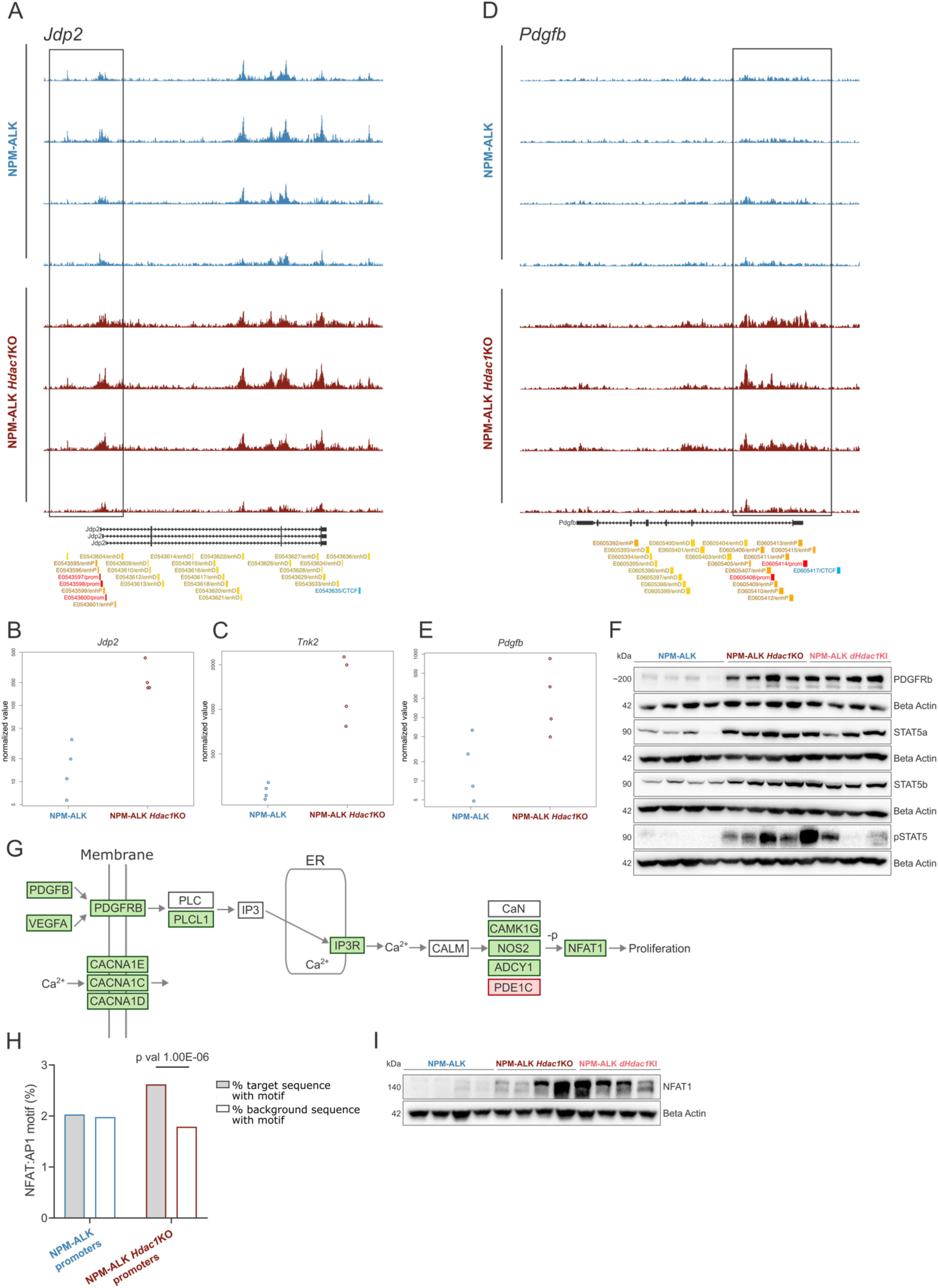
Loss of *Hdac1* hyperactivates oncogenic transcription. (A) ATAC-seq tracks downloaded from the UCSC Genome Browser^2^ depicting peaks, which represent chromatin accessibility in the *Jdp2* gene. Biological replicates of NPM-ALK end-stage thymic tumors (n=4, blue) and of NPM-ALK *Hdac1*KO end-stage thymic tumors (n=4, red) are shown. The Gencode track (Gencode VM23 release) is displayed below the ATAC-seq tracks, indicating different transcripts of the *Jdp2* gene. Colored boxes on the bottom show ENCODE Candidate Cis-Regulatory Elements (cCREs) combined from all available cell types (red promoter, orange proximal enhancer, yellow distal enhancer, blue CTCF binding sites). (B) Normalized value for size factor for raw count visualization based on RNA-seq analysis for *Jdp2* comparing NPM-ALK (blue) and NPM-ALK *Hdac1*KO (red) samples. (C) Normalized value for size factor for raw count visualization based on RNA-seq analysis for *Tnk2* comparing NPM-ALK (blue) and NPM-ALK *Hdac1*KO (red) samples. (D) ATAC-seq tracks downloaded from the UCSC Genome Browser^2^ depicting peaks, which represent chromatin accessibility in the *Pdgfb* gene. Biological replicates of NPM-ALK end-stage thymic tumors (n=4, blue) and of NPM-ALK *Hdac1*KO end-stage thymic tumors (n=4, red) are shown. The Gencode track (Gencode VM23 release) is displayed below the ATAC-seq tracks, indicating the *Pdgfb* transcript. Colored boxes on the bottom show cCREs as in (A). (E) Normalized value for size factor for raw count visualization based on RNA-seq analysis for *Pdgfb* comparing NPM-ALK (blue) and NPM-ALK *Hdac1*KO (red) samples. (F) Western blot showing protein levels of PDGFRb, STAT5a, STAT5b and pSTAT5 in end-stage thymic tumors excised from NPM-ALK (n=4), NPM-ALK *Hdac1*KO (n=4) and NPM-ALK *Hdac1*KI mice (n=4). Beta-Actin was used as a loading control. Numbers on the left indicate the molecular weight of analyzed proteins in kiloDalton (kDa). (G) Schematic representation of Ca^2+^ signaling in a cell. Green boxes indicate upregulation in NPM-ALK *Hdac1*KO tumors as compared to NPM-ALK tumors (RNA-seq) and red boxes indicate downregulation in NPM-ALK *Hdac1*KO tumors as compared to NPM-ALK tumors (RNA-seq). ER stands for *endoplasmic reticulum*. (H) Bar plot depicting the results of the Homer Motif analysis^3^, indicating enrichment of the NFAT:AP1 motif in promoter peaks of NPM-ALK *Hdac1*KO samples (red) compared to NPM-ALK samples (blue) or control sequences (background) identified by ATAC-seq analysis. (I) Western blot showing protein levels of NFAT1 in end-stage thymic tumors excised from NPM-ALK (n=4), NPM-ALK *Hdac1*KO (n=4) and NPM-ALK *Hdac1*KI mice (n=4). Beta-Actin was used as a loading control. Numbers on the left indicate the molecular weight of analyzed proteins in kiloDalton (kDa).

Importantly, we found an upregulation of the PDGFRB-STAT5-IL10 oncogenic axis, which was recently shown to be crucial for the aggressiveness of ALK+ ALCL^56^. In NPM-ALK *Hdac1*KO tumors, chromatin accessibility in the promoter region of the *Pdgfb* gene was highly increased (LFC = 9.4, p = 1.69E-4) (**Figure 6D**), concomitant with a significant upregulation of *Pdgfb* mRNA (LFC = 3.69, adj p = 1.80E-02) (**Figure 6E**). Moreover, increased gene expression of *Vegfa, Pdgfrb, Stat5a* and *Il10* was observed in NPM-ALK *Hdac1*KO tumors (|LFC| ≥ 1, adj p < 0.05) (**Supplementary Figure 6D**). The hyperactivation of the PDGFRB-STAT5 axis was corroborated at the protein level in biological replicates of NPM-ALK *Hdac1*KO tumors, demonstrating consistent upregulation of PDGFRB, STAT5A/B and phosphorylation of total STAT5 (**Figure 6F**). Moreover, increased ALK activity and upregulation of the PDGFRB- STAT5-IL10 oncogenic axis were confirmed in NPM-ALK *Hdac1*KI tumors, showcasing that their deregulation likely depends on catalytic activity of HDAC1.

The PDGFRB is implicated in multiple pathways and its activation can also lead to the release of Ca^2+^ from the *endoplasmic reticulum* (ER). Furthermore, it was shown that NPM-ALK can mimic TCR signaling, mostly *via* the oncogenic Ras pathway, but it is also weakly coupled to the calcium/NFAT pathway^57^. Notably, calcium signaling was among the top significantly enriched pathways identified in NPM-ALK *Hdac1*KO tumors based on gene expression data (**Figure 5G**, **Supplemental Table 1)**. Several components of the calcium pathway, including *Plcl1*, *Itpr3*, *Camk1g*, *Nos2*, *Adcy1,* as well as the TF *Nfat1* were significantly upregulated (|LFC| ≥ 1, adj p < 0.05) (**Figure 6G**, **Supplementary Figure 6E**). Calcium dependent NFAT TFs can act synergistically with AP-1 TFs^58^, which are known to be aberrantly expressed in ALK+ ALCL^59^. Accordingly, we further examined TF motifs in open chromatin regions in NPM-ALK and NPM-ALK *Hdac1*KO tumors. The NFAT:AP1 motif was significantly overrepresented (p = 1.00E-06) compared to background in NPM-ALK *Hdac1*KO tumors but not in NPM-ALK tumors (**Figure 6H**). NFAT proteins are furthermore key regulators of T cell development^60^ and could potentially help explain the deregulation of T cell-specific pathways as seen in the transcriptomics analyses (**Figure 5G**). Upregulation of NFAT1 in NPM-ALK *Hdac1*KO tumors as well as in NPM-ALK *Hdac1*KI tumors was further confirmed on protein level (**Figure 6I**). Upon prolonged Ca^2+^ signaling, ER Ca^2+^ can become depleted and extracellular Ca^2+^ influx is initiated to maintain the signaling. Along these lines, we observed a significant upregulation of the calcium channel encoding genes *Cacna1e, Cacna1c* and *Cancna1d* in NPM-ALK *Hdac1*KO tumors (|LFC| ≥ 1, adj p < 0.05) (**Figure 6G**, **Supplementary Figure 6F**). All in all, loss of *Hdac1* in T cells results in hyperactivation of pro-oncogenic transcription programs, suggesting that the observed accelerated lymphomagenesis is likely a consequence of synergistic effects of multiple deregulated pathways, with a strong involvement of PDGRFB- and Ca^2+^ signaling. Moreover, the accelerated lymphomagenesis and deregulation of oncogenic pathways in NPM-ALK-transgenic mice was highly dependent on the catalytic activity of HDAC1, since the same pathways were consistently found to be deregulated in NPM-ALK *Hdac1*KI tumors.

## DISCUSSION

Our study contributes novel insights into the intricate roles of HDACs in the context of T cell lymphoma. We showed that T cell-specific deletion of *Hdac1* or *Hdac2* drastically accelerates NPM-ALK driven lymphomagenesis, with a notably more pronounced effect observed upon HDAC1 loss. This finding suggests a distinct contribution of HDAC1 and HDAC2 to the transformation process of T cells, despite their high sequence homology. Interestingly, pharmacological inhibition of HDACs using Entinostat yielded contrasting results compared to genetic loss of HDAC1 protein or enzymatic activity. Entinostat treatment significantly delayed or even prevented tumor development in pre-tumorigenic mice, despite the persistent activity of NPM-ALK signaling. This discrepancy needs to be further evaluated, but might be explained by the following reasons. Firstly, Entinostat as a class I specific HDACi inhibits HDAC1 as well as HDAC2 and HDAC3, while in the case of genetic loss of *Hdac1*, HDAC2 and HDAC3 remain expressed. Similarly, complete loss of HDAC1 and HDAC2 in thymocytes results in a block in T cell development, while gradual loss of HDAC activity induces lymphoblastic lymphoma^19^. Furthermore, the observed acute thymic involution following Entinostat treatment, also described in nonclinical safety assessment of another HDACi Vorinostat^61^, raises questions about the systemic effects of HDAC inhibition on the tumor microenvironment and immune cell compartments. We speculate that changes in T cell progenitors in the bone marrow may contribute to the observed phenotype, suggesting a broader impact of HDAC inhibition beyond tumor cells alone. Moreover, the fact that prolonged effects of HDACi were observed months after cessation of the 2-week treatment of young mice, suggests that the treatment eradicated a transient early developmental progenitor cell or even early transformed lymphoma stem cells^62^, which would normally give rise to NPM-ALK lymphoma. This might also be supported by the frequent incidence of ALCL in children and young adults^63^.

Our results challenge the conventional paradigm of HDACs primarily functioning as transcriptional repressors. Surprisingly, deletion of *Hdac1* did not lead to the anticipated stochastic genome-wide chromatin opening, but rather resulted in both transcriptional repression and upregulation of gene expression. Our findings support a model wherein HDACs, in conjunction with HATs, play a crucial role in maintaining the delicate balance of histone acetylation patterns, thereby dynamically regulating gene transcription^64–66^.

Some of the observed effects might also stem from indirect consequences of HDAC depletion, such as activation of transcriptional activators or loss of repressive factors, which would result in transcriptional activation of target genes. The upregulation of the calcium-dependent TF NFAT1 and induction of its downstream targets in our model might represent such a mechanism. Additionally, HDACs target non-histone proteins and changes in overall protein acetylation might contribute to the observed phenotype. Indeed, in a previous study we identified several hundred differentially acetylated proteins, including chromatin modifying proteins and transcription factors, in mouse NPM-ALK tumor cell lines depleted for HDAC1, using quantitative acetylomics^15^.

The loss of *Hdac1* selectively perturbed cell type-specific transcription, in line with previous studies demonstrating the essential function of HDACs to maintain lineage-specific gene expression^50,51^. Notably, depletion of HDAC1 protein or loss of its catalytic activity resulted in significant alterations of the immunophenotype of ALK+ tumor cells, including a higher number of TCRb expressing cells and consequently increased dissemination of tumor cells into distant organs. The switch in immunophenotype and the hyperactivated oncogenic signaling including the PDGFRB-STAT5-IL10 oncogenic axis might suggest epigenetic reprogramming of ALK+ tumor cells upon loss of HDAC1. Alternatively, the perturbed T-cell development might have resulted in the transformation of a different T cell lineage in *Hdac1* KO thymocytes. The latter would underscore the potential of NPM-ALK to transform a variety of T cell subtypes, which is reflected in controversial findings regarding the cell or origin of ALK+ ALCL^47,67–71^.

In conclusion, our study sheds light on the intricate roles of HDAC1 and HDAC2 in ALCL development and highlights the therapeutic potential of HDAC inhibitors, such as Entinostat, in this context. Further elucidation of the underlying mechanisms and exploration of combinatorial therapeutic approaches are warranted to optimize treatment strategies for ALCL and other hematological malignancies.

## ACKNOWLEDGMENTS

This work was supported by a grant from the Austrian Science Fund (FWF) (project no.: I 4066). CUPM was supported by the Austrian Science Fund (FWF) (project no.: 32771). KD was supported by Austrian Science Foundation (FWF) SFB F83. MZ is a PhD candidate at Medical University of Vienna. This work is submitted in partial fulfillment of the requirement for the PhD.

We would like to thank Michaela Schlederer for her expertise and help with IHC stainings, Melanie R Hassler, Elisa Redl and Alexandra Zisser for initial help in setting up the mouse models, Astrid Haase for help in the lab, Andrea Alvarez Hernandez for staining the human TMA, Prof. Iris Gratz for fruitful discussions and finally Thomas Krausgruber, Prof. Christoph Bock and CeMM sequencing facility for sequencing and expertise in ATAC-seq data analysis.

## AUTHORSHIP

### Contribution

MZ, KD, VD, SW, RFS, KM, HF, HS and AIS performed the experiments; MZ, KD, VD and SW performed the mouse experiments; AM, CUPM and RST performed bioinformatics analysis; AIS and CS provided materials; MZ analyzed the results and made the figures; GE conceptualized the project; GE acquired funding; WE, CS and GE supervised; GE and MZ conceived the experiments and MZ wrote the manuscript.

