## Supplementary material for "HDAC1 acts as tumor suppressor in ALK-positive anaplastic large-cell lymphoma: Implications for HDAC inhibitor therapy"

#### SUPPLEMENTARY METHODS

##### Human TMA – further information

The IHC stainings were performed as sequential double-stainings, using HDAC1- and HDAC2- specific antibodies in the first round, CD3- and CD30-specific antibodies in the second round, respectively. TMAs containing ALK+ and ALK- ALCL samples were stained for HDAC1/2 and CD30, while TMAs containing PTCL and AITL samples were stained for HDAC1/2 and CD3 protein expression.

First Round detection: BOND Polymere Refine Detection, Leica Biosystems DS9800

Second Round detection: BOND Polymere Refine Red Detection, Leica Biosystems DS9390

Antibodies used:

CD3 (Leica Biosystems NCL-L-CD3-565), 1:100 heat-induced epitope retrieval (HIER) 20min Epitope Retrieval Sol.1, Leica Biosystems AR9961

CD30 (DAKO M0751), 1:200 HIER 20min Epitope Retrieval Sol.2, Leica Biosystems AR9640

HDAC1 (kind gift of Christian Seiser, clone 10E2), 1:200 HIER 20min Epitope Retrieval Sol.2, Leica Biosystems AR9640

HDAC2 (kind gift of Christian Seiser, clone 3F3), 1:200 HIER 20min Epitope Retrieval Sol.2, Leica Biosystems AR9640

**Mice genotyping**

All mice were genotyped twice, after weaning and after exitus. DNA was isolated with DirectPCR Lysis Reagent (Tail) (Viagen, cat#102-T) from ear or tail tissue. Genotyping was performed with GoTaq® Green Master Mix (Promega) according to the manufacturer's suggestions. The PCR products were visualized on 2% agarose gels containing 0.01 $\mu$L/mL Midori Green (Nippon Genetics Europe #MG04).

Genotyping was performed with the following primers:

ALK forward: 5'-GGTTCAGGGCCAGTGCATAT-3'

ALK reverse: 5'-CTGGCCTTCATACACCTCCC-3'

CRE forward: 5'-ATGCTTCTGTCCGTTTGCCG-3'

CRE reverse: 5'-TGAGTGAACGAACCTGGTCG-3'

Hdac1 forward: 5'-GGTAGTTCACAGCATAGTACTT-3'

Hdac1 reverse: 5'-CCTGTGTCATTAGAATCTACTT-3'

Hdac2 forward: 5'-GGTAGTTCACAGCATAGTACTT-3'

Hdac2 reverse: 5'-GTTACGTCAATGACATCGTCTT-3'

Hdac1 KI screen forward: 5'-GCATCGCCTTCTATCGCCTTC-3'

Hdac1 KI screen reverse: 5'-CTTGGTCATCTCCTCAGCATTGG-3'

Rosa26 screen forward: 5'-AAGAACTGCAGTGTTGAGGC-3'

Rosa26 screen reverse: 5'-TCTCCCAAAGTCGCTCTGAG-3'

#### Immunohistochemistry (IHC)

After fixation in 4.5 % neutral buffered formaldehyde solution tissues were embedded in paraffin and cut into 5 µM sections. Following deparaffinization and rehydration, antigen retrieval was performed by heat-treatment in TRIS/EDTA (Target Retrieval Solution pH 9 Dako) or citrate buffer (Target Retrieval Solution pH 6 Dako). Endogenous peroxidase was blocked by 3% $H_2O_2$, slides were blocked with avidin/biotin (Vector lab), super block (Empire Genomics) and universal mouse block (Empire Genomics). Slides were incubated with primary antibody o/n at 4°C, followed by secondary antibody and incubation with the IDetect™ Super Stain System - HRP kit (Empire Genomics). The staining was performed with AEC reagent (Empire Genomics), followed by counterstain by incubation in Mayer's hemalum solution (Merck).

Antibodies used:

Ki67 (D3B5) Cell Signaling #9129 1:400 dilution

CC3 (ASP175) Cell Signaling #9661 1:200 dilution

ALK (D5F3) Cell Signaling #3633 1:250 dilution

#### Protein isolation, Western blotting – buffers composition and antibodies

Buffer compositions:

Hunt buffer: 20 mM Tris pH 8, 100 mM NaCl, 1 mM EDTA, 0.5% NP-40, protease inhibitor - Roche

Milk blocking solution: 5% skim milk powder, 1% PVP, 0.01% Sodium azide in TBS-T (=TBS containing 1% Triton-X100)

BSA blocking solution: 5% BSA, 0.01% Sodium azide in TBS-T (=TBS containing 1% Triton-X100)

*BSA blocking solution was used when phosphorylated proteins were detected.

ECL solutions: Amersham ECL Detection Reagents or Bio-Rad Clarity ECL substrate (to detect pSTAT5)

Antibodies used:

HDAC1 (kind gift of Christian Seiser, clone 10E2), dilution 1:1000

HDAC2 (kind gift of Christian Seiser, clone 3F3) dilution 1:1000

ALK Cell Signaling #4691, dilution 1:2500

p-ALK Cell Signaling #6941, dilution 1:500

STAT3 Cell Signaling #12640, dilution 1:1000

p-STAT3 Cell Signaling #9131, dilution 1:1000

CD3g Proteintech #21120-AP, dilution 1:1000

CD3d Proteintech #16669-I-AP, dilution 1:1000

CD3e (SP7) Abcam #ab16669, dilution 1:1000

PDGFRb Cell Signaling #3169, dilution 1:1000

STAT5a (C-6) Santa Cruz #sc-271542, dilution 1:10000

STAT5b (G-2) Santa Cruz #sc-1656, dilution 1:1000

pSTAT5 Cell Signaling #9351, dilution 1:1000

NFAT1 Cell Signaling #4389, dilution 1:1000

c-MYC (E5Q6W9) Cell Signaling #18583, dilution 1:500

Beta Actin (Proteintech #66009-I-Ig), dilution 1:5000

Alpha Tubulin (Proteintech #66031-1-Ig), dilution 1:5000

Goat anti-rabbit horseradish peroxidase-linked secondary antibody (Jackson Immuno Research, #111-036-047), dilution 1:10,000

Rabbit anti-mouse horseradish peroxidase-linked secondary antibody (Jackson Immuno Research, #315-035-045), dilution 1:10,000

**ATAC-seq and RNA-seq sample preparation – further information and buffer compositions**

Snap frozen tumor tissue was lyzed in homogenization buffer. Nuclei were pelleted by centrifugation (5 min, 4°C, 350 rcf). The supernatant was used to isolate RNA (see below). Nuclei were resuspended in homogenization buffer and gradients of iodixanol solution were used to obtain the band containing the nuclei at the 30 – 40% iodixanol interface after centrifugation (20 min, 4°C, 3000 rcf, break off). Nuclei were diluted in ATAC-RSB-Tween buffer and counted. 50,000 nuclei per sample were centrifuged (10 min, 500 rcf, 4°C) and resuspended in ATAC-seq Reaction Mix. Reactions were incubated (37°C, 30 min, 1000 rpm). After incubation, Binding Buffer (Qiagen MinElute PCR Purification kit) was added. A cleaning protocol using the MiniElute PCR Purification kit was performed according to the manufacturer and samples were eluted in Elution Buffer.

RNA isolation: 150 $\mu$L of the supernatant (see above) was mixed with Trizol and chloroform, vortexed and centrifuged (15 min, 4°C, 21,000 rcf). The aqueous layer was mixed with an equal volume of 100% ethanol and passed through a QIAgen RNeasy column. The QIAgen RNeasy protocol was followed and samples were eluted in 27 μL Elution Buffer. 3 μL of 10 x Turbo™ DNase Buffer and 1 μL of Turbo™ Dnase enzyme was added and the samples were incubated (30 min, 37°C). 70 μL of RNase-free water and 350 μL of QIAgen RLT Buffer were added. 250 μL of 100% ethanol was added and the samples were applied to the column and washed 2 x with RPE. Samples were eluted in RNase-free water.

Buffer compositions:

Homogenization buffer: final concentration 0.26 M sucrose, 0.03 M KCl, 0.01 M $MgCl_2$, 0.02 M Tricine-KOH pH 7.8, with 0.001 M DTT, 0.5 mM spermidine, 0.15 mM spermine, 0.3 % NP40, 1x cOmplete™ Protease Inhibitor, 10 μL RiboLock per mL of buffer

Iodixanol solution: iodixanol diluted with appropriate amount of diluent buffer

Diluent buffer: 0.15 M KCl, 0.03 M $MgCl_2$, 0.12 M Tricine-KOH pH 7.8

ATAC-RSB-Tween buffer: final concentration 0.01 M Tris-HCl pH 7.5, 0.01 M NaCl, 0.003 M $MgCl_2$, 0.1 % Tween-20 in water

ATAC-seq Reaction Mix per sample: 5 μL water, 16.5 μL PBS, 25 μL 2x TD, 0.5 μL 1% digitonin, 0.5 μL 10% Tween-20, 2.5 μL Tn5

**RNA-seq – additional information**

Cutadapt[1] was used to remove unwanted sequences (e.g., adapters, poly-A tails, etc.) and low-quality reads on the FASTQ files provided by the sequencing facility. FASTQC[2] and MULTIQC[3] were both used to check the quality of reads. The STAR aligner tool[4] was used to map the reads to the GRCm39 mouse reference genome, obtained from ENSEMBL (https://www.ensembl.org/Mus_musculus/Info/Index).

Htseq-count[5] was used to obtain gene counts. Finally, we performed a naive pre-filtering step to remove low count genes by only keeping those transcripts with an average count across all samples bigger than one. We used the R/Bioconductor package DESeq2[6] to perform differential gene expression (DE) analysis of the Htseq transcript counts. Genes with an adjusted P-value (p adj) $< 0.05$ and absolute log2 fold change (logFC)$\geq 1$ were considered significantly differentially expressed. Finally, read counts were normalized by the DESeq2 normalization method of variance stabilizing transformation (VST), to be used primarily for visualization. Figures for this analysis were generated using DESeq2.

**ATAC-seq – further information**

Initial data quality check was done using FastQC[2]. Next, transposase (Tn5) sequences were removed using the cutadapt tool[1]. All Illumina adapters were likewise removed. Next Bowite2[7] was used for mapping reads to the reference genome. Sorting of the mapped reads was done using samtools[8]. Mitochondrial reads and sex chromosomes as well as reads with low mapping quality (MAPQ $< 30$) were removed. For deduplication rmdup and samtools[8,9] were used. Peaks were called with MACS2 (Model-based analysis of ChIP-seq)[10]. FRiP scores were calculated and quality check was performed using multiQC[3]. Counting reads per peak was accomplished with featureCounts[11]. To annotate peaks, the package ChIPseeker[12] was used. Differential enrichment analysis was done using edgeR[13]. Motif analysis was done using HOMER software[14].

**Functional gene set enrichment analysis**

Ingenuity Pathway Analysis (IPA®,Qiagen) was performed including deregulated genes (RNA-seq: p adj < 0.05 and logFC ≥ I1I), which showed a correlation with changes in chromatin accessibility (p for correlation < 0.05).

**HDAC activity assay**

20 µg/20 µL of protein were incubated with 4 µL of $^{3}H$-acetate–labelled chicken erythrocyte histone mix (1.5 mg/mL) for 1 h at 30°C on a thermoshaker at 300 rpm. The reaction was stopped by adding 35 µL of histone stop solution and 800 µL ethyl acetate. The samples were vortexed and centrifuged (4 min, 10000 rpm). 600 µL of the organic phase was transferred to 3 mL of scintillation solution and mixed gently. The HDAC activity was determined by a Liquid Scintillation Analyzer.

**FACS Immunophenotyping – preprocessing of the samples**

Single cell suspensions of thymus, tumor and spleen were obtained by passaging the tissues through a 70 µm nylon cell strainer in staining buffer (PBS supplemented with 2% FCS). Bone marrow was isolated using empty DMEM medium following the STAR protocol[15]. Erythrocytes were lysed (5 min on ice) using erylysis buffer (final concentrations 0.15 M NH4Cl, 10 mM KHCO3, 0.1 mM EDTA, pH = 7.2 – 7.4) before staining.

**FACS Immunophenotyping – antibodies**

Antibodies used:

| ANTIGEN | FLUOROPHORE | CLONE | COMPANY |
|---|---|---|---|
| viability | Zombie NIR | / | BioLegend |
| CD45 | BV650 | 30-F11 | BD Biosciences |
| CD19 | APC-Cy7 | 6D5 | BioLegend |
| CD11b | PerCP-Cy5.5 | M1/70 | Invitrogen Antibodies |
| Gr-1 | PE | RB6-8C5 | Invitrogen Antibodies |
| CD117 = cKit | PECY5 | 2B8 | Invitrogen Antibodies |
| Sca1 | PE-Cy7 | D7 | Invitrogen Antibodies |
| CD34 | AF700 | RAM34 | Invitrogen Antibodies |
| CD16/32 | PerCP-eF710 | 93 | Invitrogen Antibodies |
| CD127 = IL7R | FITC | A7R34 | Invitrogen Antibodies |
| TCRb | BV510 | H57-597 | BD Biosciences |
| TCRgd | BV711 | GL3 | BD Biosciences |
| CD8a | BUV496 | 53-6.7 | BD Biosciences |
| CD4 | BUV395 | GK1.5 | BD Biosciences |
| CD25 | PB | PC61 | BioLegend |
| CD44 | BV570 | IM7 | BioLegend |
| ALK (cell signaling#3633) + AF647 goat anti-rabbit (Invitrogen#a-21244) | AF647 | / | Invitrogen |
| CD69 | BV785 | H1.2F3 | BioLegend |
| CD62L | BV605 | MEL-14 | BD Biosciences |

**FACS Immunophenotyping – tSNE**

For unsupervised clustering, first the thymic tumor samples were downsized using FlowJo™ (BD Life Sciences) plugin "Downsample" (version 3.3.1). 100.000 of events were used from the gating for alive leukocytes (CD45+ cells, negative for viability stain). New Downsample populations were concatenated (using all events and all compensated parameters). Next tSNE analysis (part of FlowJo™, BD Life Sciences) was performed on concatenated sample. To run tSNE, all markers were used except viability and CD45.

**In vivo treatments – further information**

Mice were treated with HDACi for two consecutive weeks on a five-days-on-two-days-off schedule. Entinostat (Selleckchem) was administered via interperitoneal (IP) injection, with a 1 mL single use Tuberculin syringe (Chirana) and a 27-gauge (G) x ¾” (0.4 mm x 20 mm) safety needle (SOL-CARE). Entinostat (Selleckchem) was aliquoted beforehand and diluted daily prior to injection. The drug was diluted in 90% sterile filtered corn oil (Mazola) and 10% DMSO. Dilutions for different treatment doses can be seen in Table 1. The volumes injected based on mouse weight are shown in Table 2 (example for 10 *μ*g/g treatment dose).

Table 1

| Treatment group | mg of substance/mL DMSO+oil (10/90) |
|---|---|
| Entinostat 50 *μ*g/g | 10 mg/mL |
| Entinostat 20 *μ*g/g | 4 mg/mL |
| Entinostat 10 *μ*g/g | 2 mg/mL |
| Entinostat 5 *μ*g/g | 1 mg/mL |

Table 2

| Body weight (g) | Dose (*μ*g) | Vol. injection (*μ*L) |
|---|---|---|
| 15 | 150 | 75 |
| 16 | 160 | 80 |
| 17 | 170 | 85 |
| 18 | 180 | 90 |
| 19 | 190 | 95 |
| 20 | 200 | 100 |
| 21 | 210 | 105 |
| 22 | 220 | 110 |
| 23 | 230 | 115 |
| 24 | 240 | 120 |
| 25 | 250 | 125 |

**Electron Microscopy**

After fixation in Karnovsky solution, tissues were postfixed in 1% osmium ferrohexacyanoferrate II, then washed in distillated water and dehydrated in ethanol and propylene oxide and embedded in Epon resin (Embed 812). Ultrathin sections of 70 nm were made on a Leica-Ultracut- EM-UC7. For contrast enhancement, the ultrathin sections were stained in 2% uranyl acetate and 1% lead citrate. Transmission electron microscopy was then performed on a TEM Jeol 1400 Plus (Jeol) at 60kV, pictures were taken with Quemesa_Camera in radius-software.

**Statistics**

Survival statistics were analyzed using GraphPad Prism (version 8.4.3). To assess differences between groups pairwise curve comparison using Log-rank (Mantel–Cox) test was performed.

Data are represented as mean ± SD, if not otherwise specified and were analyzed using GraphPad Prism (version 8.4.3). To assess differences between groups, unpaired t test or one-way ANOVA were used, depending on number of sample groups.

Significance was defined according to following P-values: *P < 0.05; **P < 0.01; ***P < 0.001 ****P < 0.0001.

#### SUPPLEMENTARY FIGURES CAPTIONS

##### SUPPLEMENTARY FIGURE 2: *Hdac1* loss in T cells accelerates lymphomagenesis

(A) Kaplan Meier survival analysis of NPM-ALK mice (n=25, light blue line), *Hdac1*KO mice (n=25, light gray line) and *Hdac2*KO mice (n=17, dark gray line) in biological replicates. GraphPad Prism version 8.4.3 was used for analysis.

(B) Representative microscopic images of liver sections from NPM-ALK mice (n=2) stained for ALK expression by IHC. Sections were counterstained with hematoxylin (blue). Infiltration of ALK positive cells around vessels is displayed at two magnifications. Scale bar represents 100 μm (upper panel), and 50 μm (lower panel).

##### SUPPLEMENTARY FIGURE 3: Accelerated lymphomagenesis depends on HDAC1 enzymatic activity

(A) Kaplan Meier survival analysis of NPM-ALK mice (n=25, blue line), *Hdac1*KO mice (n=25, light gray line) *dHdac1*KI mice (n=19, dark gray line) in biological replicates. GraphPad Prism version 8.4.3 was used for analysis.

(B) Immunoblot of protein levels of HDAC1 and HDAC2 in end-stage thymic tumors excised from NPM-ALK (n=4), NPM-ALK *Hdac1*KO (n=4) and NPM-ALK *Hdac1*KI (n=4) mice. Beta-Actin was used as a loading control. The molecular weight of analyzed proteins in kiloDaltons (kDa) is indicated by the numbers on the left. Corresponding quantification of the blots is shown in Figure 4J.

**SUPPLEMENTARY FIGURE 4: Loss of HDAC1 protein or HDAC1 catalytic activity causes changes in immunophenotype**

(A) Exemplar gates utilized for analyzing thymic tumor samples, established using WT thymus samples stained and acquired concurrently with tumor samples.

(B) Exemplar gates employed for analyzing spleen samples, established using WT spleen samples stained and acquired concurrently with samples from tumor-bearing mice.

(C) Exemplar gates applied for analyzing bone marrow samples, established using WT bone marrow samples stained and acquired concurrently with samples from tumor-bearing mice.

(D) Unsupervised clustering of flow cytometry (FACS) immunophenotyping using tSNE plots (live, CD45+ leukocytes from thymic tumors were used for clustering). From left to right, clustering of all samples (n=15) is shown, followed by NPM-ALK only (n=6), NPM-ALK *Hdac1*KO only (n=5) and NPM-ALK *dHdac1*KI only samples (n=4).

(E) Representative flow cytometry histograms showing ALK expression on live CD45+ NPM-ALK positive thymocytes, stained with either the full panel (blue) or the full panel minus ALK antibody (red), serving as the fluorescence minus one control (FMO control).

(F) tSNE plot based on CD62L expression (live, CD45+ leukocytes) of all samples (n=15). and the color depicts the expression of CD62L with red depicting high expression and green is depicting low expression. Samples as in (D).

**SUPPLEMENTARY FIGURE 5: Loss of *Hdac1* selectively perturbs cell-type specific transcription**

(A) Representative transmission electron microscopy pictures of NPM-ALK (n=1) and NPM-ALK *Hdac1*KO (n=1) end-stage thymic tumors showing the nuclear composition of hetero- and euchromatin (scale bar representing 2μm).

(B) Volcano plot depicting differential gene expression of biological replicates of NPM-ALK (n=4) and NPM-ALK *Hdac1*KO (n=4) end-stage thymic tumors based on RNA-seq analysis. Red represents significantly deregulated genes with ILFCI $\geq 1$ and adj $p < 0.05$.

(C) UCSC Genome Browser[1] ATAC-seq tracks depicting peaks, which represent open chromatin regions in *CD3d* and *Cd3g* genes. Biological replicates of NPM-ALK end-stage thymic tumors (n=4, blue) and of NPM-ALK *Hdac1*KO end-stage thymic tumors (n=4, red) are shown. Gencode tracks (Gencode VM23 release) below display corresponding transcripts. Colored boxes on the bottom show ENCODE Candidate Cis-Regulatory Elements (cCREs) combined from all available cell types (red promoter, orange proximal enhancer, yellow distal enhancer).

**SUPPLEMENTARY FIGURE 6: Loss of *Hdac1* hyperactivates oncogenic transcription**

(A) UCSC Genome Browser[1] ATAC-seq tracks depicting peaks, which represent open chromatin regions in the *Myc* gene. Biological replicates of NPM-ALK end-stage thymic tumors (n=4, blue) and of NPM-ALK *Hdac1*KO end-stage thymic tumors (n=4, red) are shown. Gencode tracks (Gencode VM23 release) below display corresponding transcripts. Colored boxes on the bottom show ENCODE Candidate Cis-Regulatory Elements (cCREs) combined from all available cell types (red promoter, orange proximal enhancer, yellow distal enhancer).

(B) Western blot showing protein expression levels of MYC in end-stage thymic tumors excised from NPM-ALK (n=4) and NPM-ALK *Hdac1*KO mice (n=4). Beta-Actin was used as a loading control. Numbers on the left indicate the molecular weight of analyzed proteins in kiloDalton (kDa).

(C) Normalized value for size factor for raw count visualization based on RNA-seq analysis for *Myc* comparing NPM-ALK (blue) and NPM-ALK *Hdac1*KO (red) samples.

(D) Normalized value for size factor for raw count visualization based on RNA-seq analysis for *Vegfa, Pdgfrb, Stat5a, Stat5b* and *Il10* comparing NPM-ALK (blue) and NPM-ALK *Hdac1*KO (red) samples.

(E) Normalized value for size factor for raw count visualization based on RNA-seq analysis for *Plcl1, Itpr3, Camk1g, Nos2, Adcy1* and *Nfat1* comparing NPM-ALK (blue) and NPM-ALK *Hdac1*KO (red) samples.

(F) Normalized value for size factor for raw count visualization based on RNA-seq analysis for *Cacna1e, Cacna1c* and *Cacna1d* comparing NPM-ALK (blue) and NPM-ALK *Hdac1*KO (red) samples.

Supplementary figure 2: *Hdac1* loss in T cells accelerates lymphomagenesis

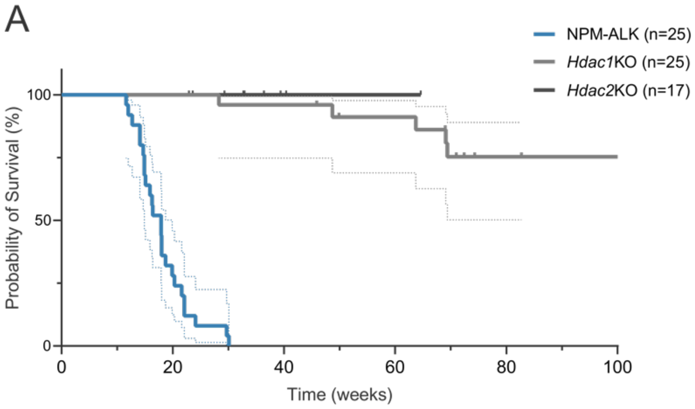


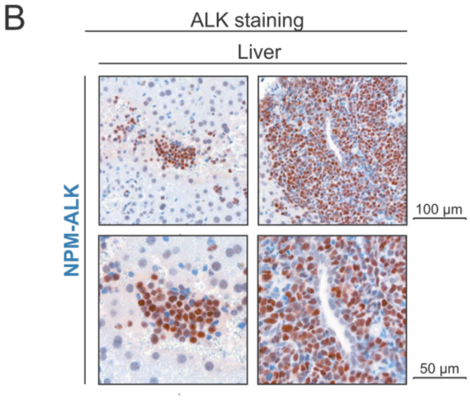

### Supplementary figure 3: Accelerated lymphomagenesis depends on HDAC1 enzymatic activity

A

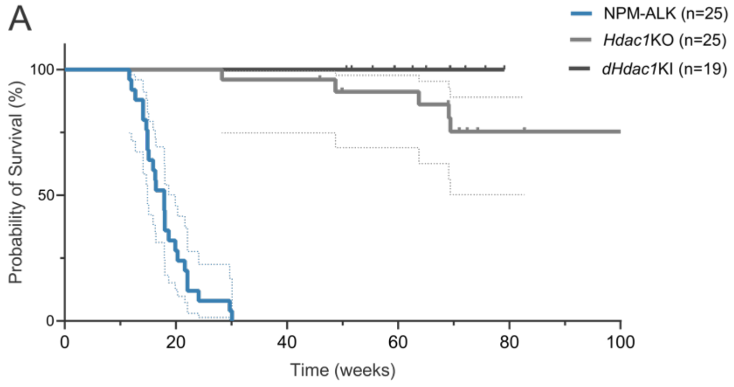


B

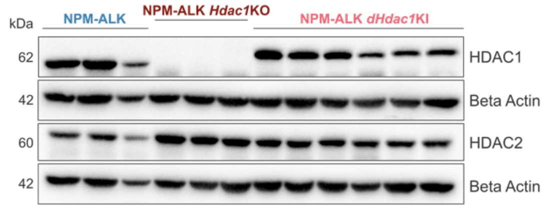

Supplementary figure 4: Loss of HDAC1 protein or HDAC1 catalytic activity causes changes in the immunophenotype

A

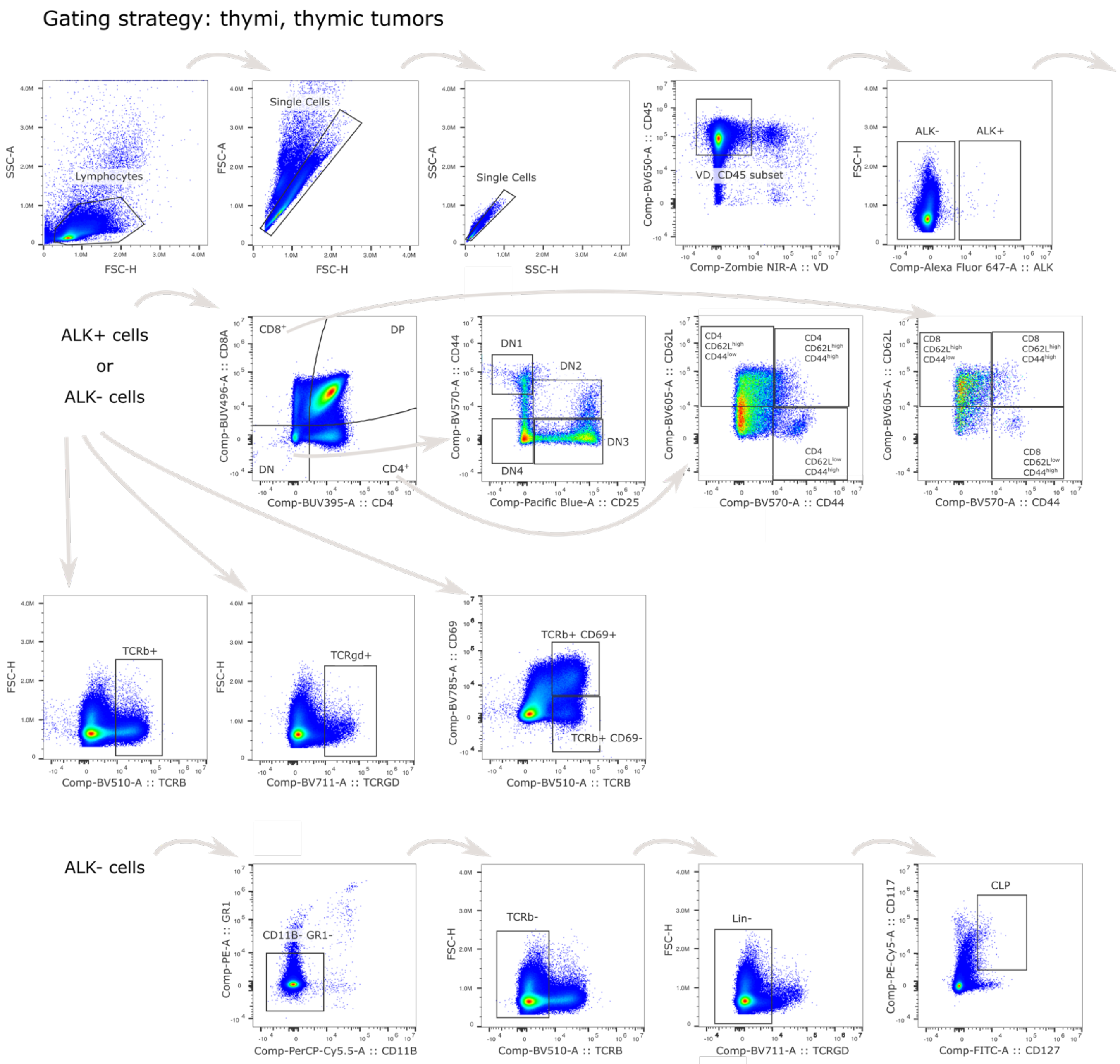

Supplementary figure 4: Loss of HDAC1 protein or HDAC1 catalytic activity causes changes in the immunophenotype

B

Gating strategy: spleen

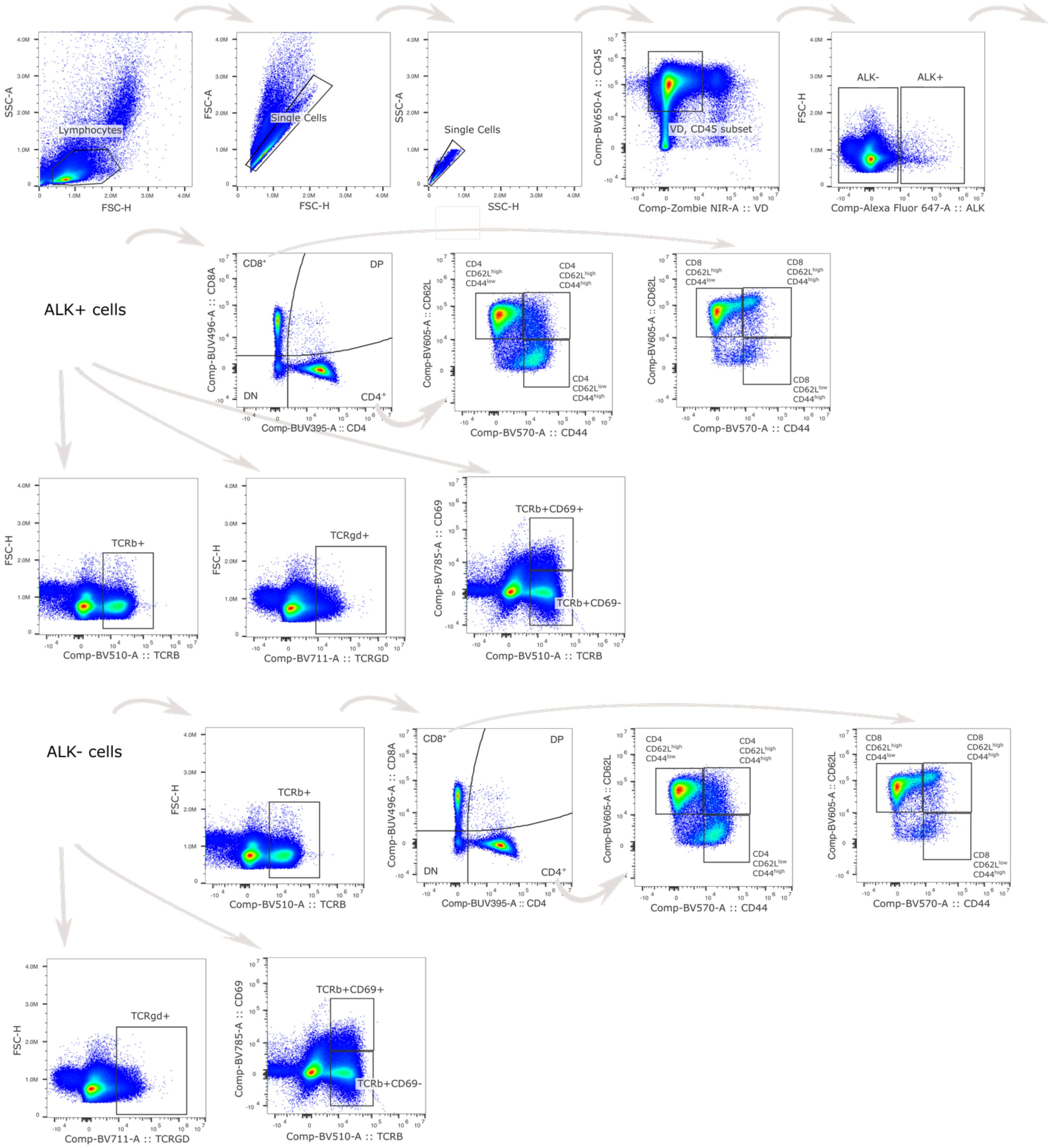

### Supplementary figure 4: Loss of HDAC1 protein or HDAC1 catalytic activity causes changes in the immunophenotype

C

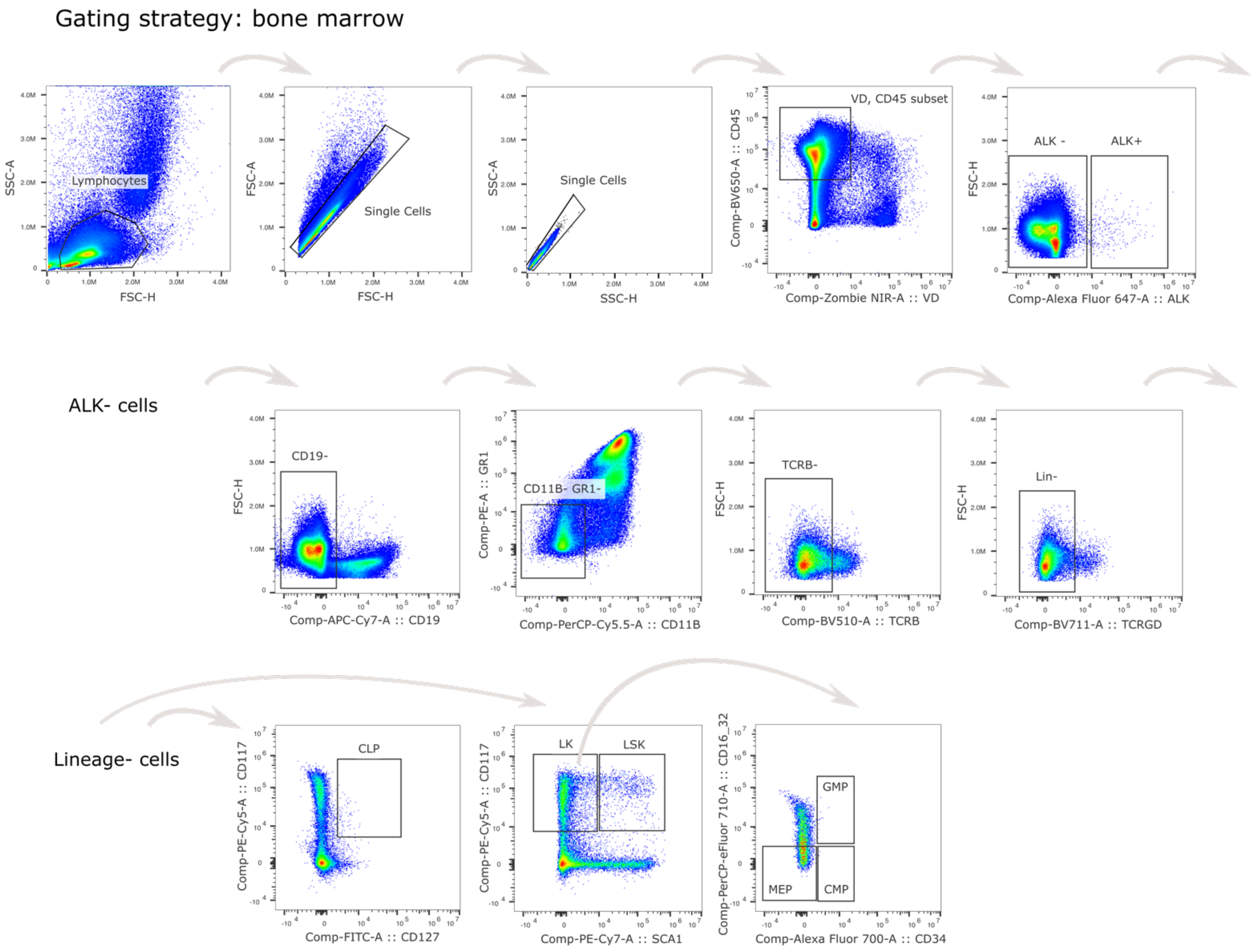


D

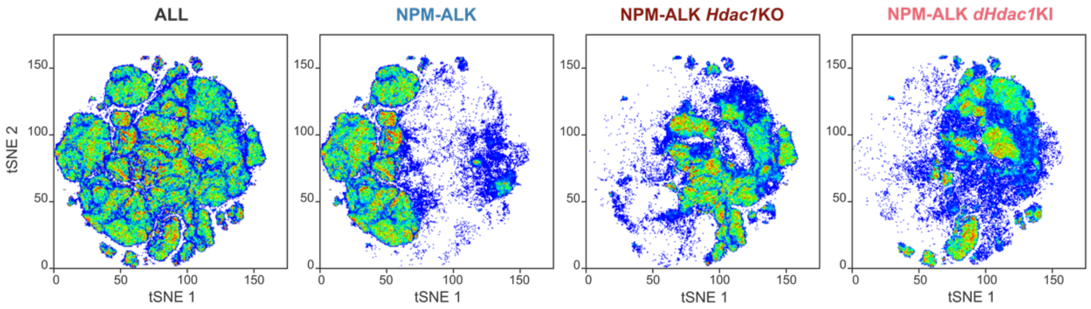


E

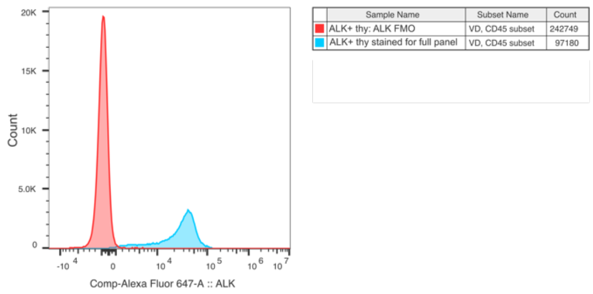


F

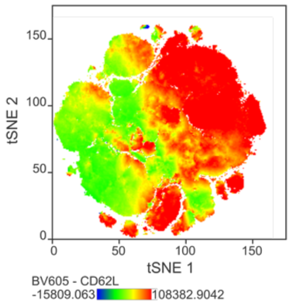

Supplementary figure 5: Loss of *Hdac1* selectively perturbs cell-type specific transcription

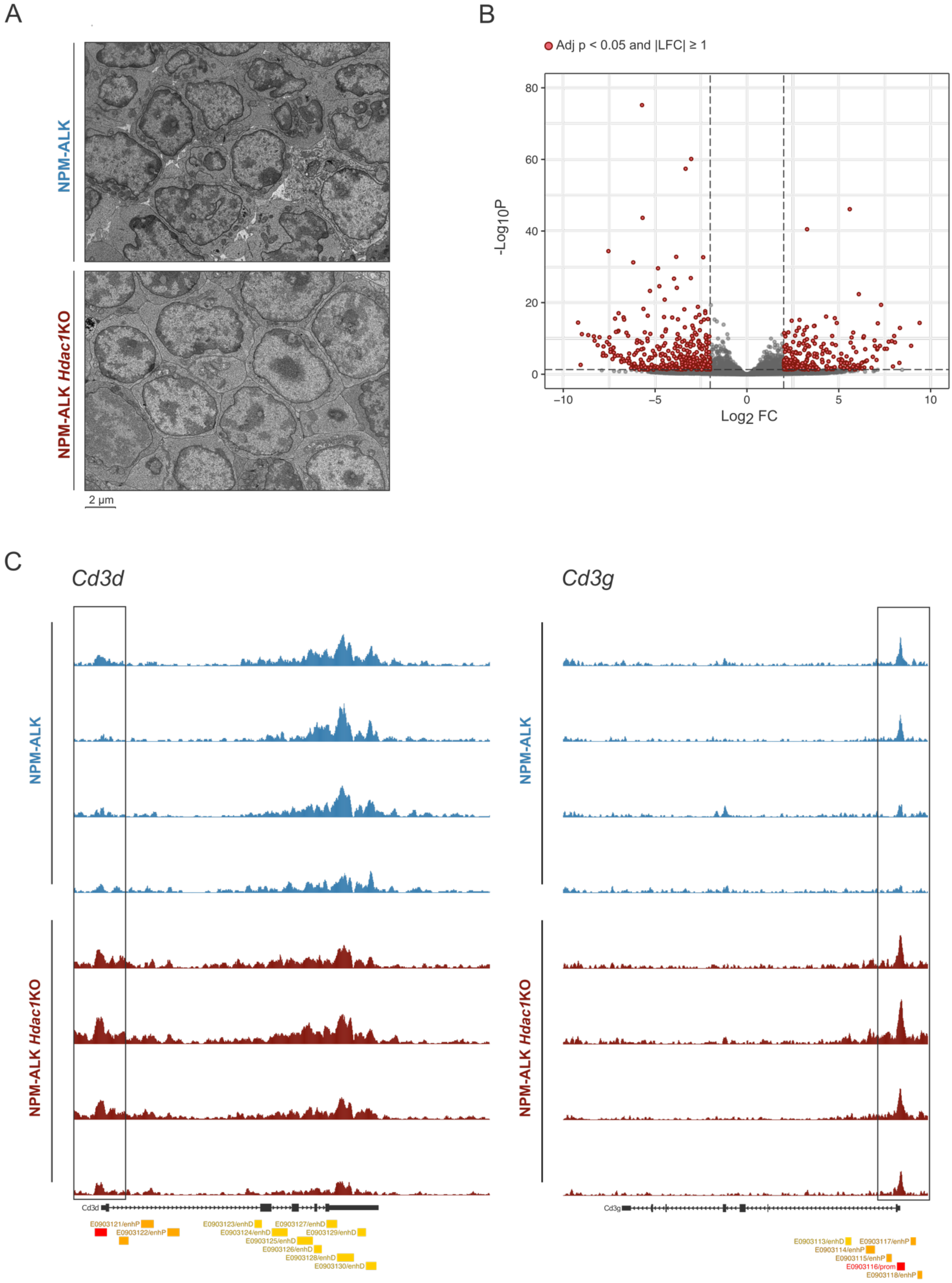

Supplementary figure 6: Loss of *Hdac1* hyperactivates oncogenic transcription

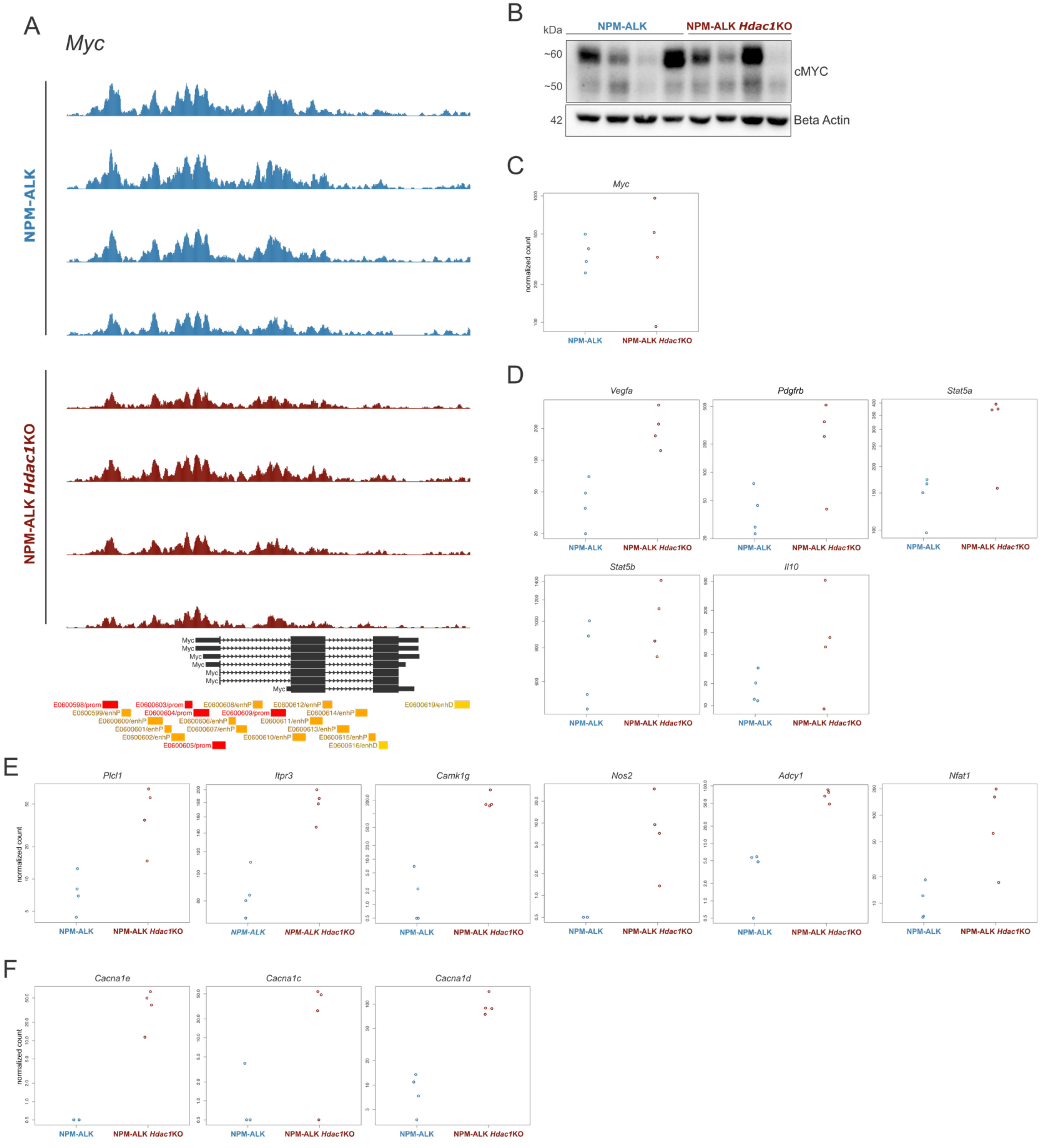

#### SUPPLEMENTARY TABLE 1

| Ingenuity Canonical Pathways | p-value | Ratio | Molecules |
|---|---|---|---|
| Calcium-induced T Lymphocyte Apoptosis | 2,04E-04 | 7,58E-02 | CD3D,CD3G,CD4,HLA-A,ITPR3 |
| OX40 Signaling Pathway | 9,55E-05 | 6,67E-02 | BCL2L1,CD3D,CD3G,CD4,HLA-A,NFKBIE |
| iCOS-iCOSL Signaling in T Helper Cells | 3,55E-05 | 6,31E-02 | CD3D,CD3G,CD4,HLA-A,ITPR3,NFKBIE,TRAT1 |
| Regulation of IL-2 Expression in Activated and Anergic T Lymphocytes | 8,51E-04 | 5,56E-02 | CD3D,CD3G,NFKBIE,TGFBR1,VAV2 |
| T Helper Cell Differentiation | 3,02E-03 | 5,48E-02 | HLA-A,TBX21,TGFBR1,TNFRSF1A |
| PKC-theta Signaling in T Lymphocytes | 4,17E-05 | 5,16E-02 | CACNA1D,CACNG4,CD3D,CD3G,CD4,HLA-A,NFKBIE,VAV2 |
| Th2 Pathway | 1,29E-04 | 5,15E-02 | CCR4,CD3D,CD3G,CD4,HLA-A,TBX21,TGFBR1 |
| CD28 Signaling in T Helper Cells | 4,79E-04 | 4,96E-02 | CD3D,CD3G,CD4,HLA-A,ITPR3,NFKBIE |
| T Cell Receptor Signaling | 1,78E-03 | 4,72E-02 | CD3D,CD3G,CD4,PAG1,VAV2 |
| Cdc42 Signaling | 1,02E-04 | 4,55E-02 | CD3D,CD3G,HLA-A,ITGA6,LIMK1,MYL10,TNK2,VAV2 |
| CTLA4 Signaling in Cytotoxic T Lymphocytes | 6,03E-03 | 4,49E-02 | CD3D,CD3G,HLA-A,TRAT1 |
| Death Receptor Signaling | 6,92E-03 | 4,35E-02 | BID,LIMK1,NFKBIE,TNFRSF1A |
| Th1 Pathway | 3,16E-03 | 4,13E-02 | CD3D,CD3G,CD4,HLA-A,TBX21 |
| Th1 and Th2 Activation Pathway | 5,25E-04 | 4,09E-02 | CCR4,CD3D,CD3G,CD4,HLA-A,TBX21,TGFBR1 |
| Apoptosis Signaling | 9,12E-03 | 4,00E-02 | BCL2L1,BID,NFKBIE,TNFRSF1A |
| Role of NFAT in Regulation of the Immune Response | 7,24E-04 | 3,87E-02 | CD3D,CD3G,CD4,HLA-A,ITPR3,NFKBIE,RCAN3 |
| PD-1, PD-L1 cancer immunotherapy pathway | 1,12E-02 | 3,77E-02 | BCL2L1,HLA-A,PDCD1,TNFRSF1A |
| PI3K Signaling in B Lymphocytes | 5,50E-03 | 3,62E-02 | CARD10,IRS2,ITPR3,NFKBIE,VAV2 |
| D-myo-inositol (1,4,5,6) / (3,4,5,6)-Tetrakisphosphate Biosynthesis | 6,17E-03 | 3,52E-02 | ALPL,DUSP10,PDCD1,PPP1R1B,PTPRF |
| 3-phosphoinositide Degradation | 9,12E-03 | 3,21E-02 | ALPL,DUSP10,PDCD1,PPP1R1B,PTPRF |
| 3-phosphoinositide Biosynthesis | 1,17E-02 | 3,01E-02 | ALPL,DUSP10,PDCD1,PPP1R1B,PTPRF |
| Calcium Signaling | 6,92E-03 | 2,91E-02 | CACNA1D,CACNG4,CAMK1G,CHRNA9,ITPR3,RCAN3 |
| T Cell Exhaustion Signaling Pathway | 1,45E-02 | 2,86E-02 | HLA-A,PDCD1,TBX21,TGFBR1,VEGFA |
| Protein Kinase A Signaling | 4,68E-04 | 2,75E-02 | ADCY1,AKAP12,DUSP10,H1-0,ITPR3,MYL10,NFKBIE,PPP1R1B,PTPRF,TCF7L1,TGFBR1 |
| Senescence Pathway | 7,59E-03 | 2,55E-02 | CACNA1D,CBX7,DHCR24,ETS2,ING1,ITPR3,TGFBR1 |
| Phospholipase C Signaling | 2,24E-02 | 2,26E-02 | ADCY1,CD3D,CD3G,ITGA6,ITPR3,MYL10 |
